# Discrete projections from MCH neurons mediate anti-parkinsonian effects of acupuncture

**DOI:** 10.1101/2023.06.07.543987

**Authors:** Ju-Young Oh, Hyowon Lee, Sun-Young Jang, Hyunjin Kim, Geunhong Park, Almas Serikov, Jae-Hwan Jang, Junyeop Kim, Seulkee Yang, Moonsun Sa, Sung Eun Lee, Young-Eun Han, Tae-Yeon Hwang, Hee Young Kim, Seung Eun Lee, Soo-Jin Oh, Jeongjin Kim, Jongpil Kim, C. Justin Lee, Min-Ho Nam, Hi-Joon Park

**Affiliations:** College of Korean Medicine, Graduate School, Kyung Hee University, Seoul 02447, Korea; Studies of Translational Acupuncture Research (STAR), Acupuncture and Meridian Science Research Center (AMSRC), Kyung Hee University, Seoul 02447, Korea; Brain Science Institute, Korea Institute of Science and Technology (KIST), Seoul 02792, Korea; Department of KHU-KIST Convergence Science and Technology, Kyung Hee University, Seoul 02447, Korea; Laboratory of Stem Cells & Cell reprogramming, Department of Chemistry, Dongguk University, Seoul 04629, Korea; Center for Cognition and Sociality, Institute for Basic Science, Daejeon 34126, Korea; Department of physiology, Yonsei University College of Medicine, Seoul, Korea; Virus Facility, Research Animal Resource Center, KIST, Seoul 02792, Korea

## Abstract

Parkinson’s disease (PD) presents with typical motor dysfunction and non-motor symptoms, including memory loss. Although acupuncture is suggested as an alternative therapy for PD, its neuroanatomical mechanisms remain unclear. We demonstrate that acupuncture ameliorates both motor and memory deficits in PD mice through activation of melanin-concentrating hormone (MCH) neurons in the lateral hypothalamus and zona incerta (LH/ZI)—MCH^LH/ZI^— via nerve conduction. We identify two distinct subpopulations of MCH^LH/ZI^ projecting to the substantia nigra and hippocampus, each of which is responsible for controlling motor and memory function. This effect can be attributed to MCH-mediated recovery from dopaminergic neurodegeneration, reactive gliosis, and impaired hippocampal synaptic plasticity. Collectively, MCH^LH/ZI^ constitutes not only the neuroanatomical basis of acupuncture but also a potential cellular target for treating both motor and non-motor PD symptoms.

**One-Sentence Summary:** Acupuncture alleviates both motor and non-motor symptoms in Parkinson’s disease by activating two distinct MCH projections.

## Main text

Current Parkinson’s disease (PD) treatments, including levodopa and deep brain stimulation (DBS), primarily address motor dysfunctions (*1*), leaving out non-motor symptoms like memory deficits. Moreover, their short-term effects, severe adverse effects, and high invasiveness have necessitated an alternative, safe, and less invasive therapeutic approach that simultaneously addresses both motor and non-motor deficits of PD. Acupuncture, a traditional East Asian medical practice for treating neurological conditions (*2, 3*), has shown potential therapeutic effects in PD patients (*4-6*) and rodent models (*7-13*). However, it is still unclear how acupuncture produces these benefits at the molecular and cellular levels.

Our previous study suggested that acupuncture’s effects on PD motor symptoms might involve the melanin-concentrating hormone (MCH) neurons (*14*), located in the lateral hypothalamus (LH) and zona incerta (ZI) (*15*). However, it is unknown whether MCH neurons project their axons to motor-related brain regions, such as the substantia nigra pars compacta (SNpc), and how they regulate the motor function. On the other hand, recent studies have shown the critical role of MCH neurons in memory processing (*16, 17*), possibly through their projections to the hippocampus (HPC) (*18, 19*). This finding motivated us to investigate the potential involvement of MCH neurons in PD-related memory loss. Therefore, our study aimed to delineate the molecular, cellular, and circuitry-level mechanisms of acupuncture’s effects on both motor and non-motor phenotypes in PD through MCH neurons.

### Acupuncture alleviates PD-like motor and memory deficits through nerve conduction

We first assessed whether the acupuncture stimulation alleviates the motor and memory deficits in the 1-Methyl-4-phenyl-1,2,3,6-tetrahydropyridine (MPTP)-induced subacute PD mouse model (*20*), which exhibits both motor and memory deficits (*21*) (Fig. 1A). Acupuncture stimulation was performed by a bilateral needle insertion into the hindlimb *Yangleungchun* (GB34; *Yanglingquan*) acupoint or a control non-acupoint to a depth of 2 mm and rotation for 30 s at a rate of two spins per second without anesthesia (Fig. 1B and 1C), as previously described (*13, 14, 22, 23*). Consistent with previous findings (*14, 22*), we found that MPTP-induced PD-like motor deficits were significantly alleviated by acupuncture at GB34, but not by non-acupoint stimulation (Fig. 1D-1J, and S1A). Additionally, by performing Y-maze and novel object recognition tests, we found that acupuncture treatment significantly alleviated MPTP-induced memory deficits, while non-acupoint stimulation did not (Fig. 1K and 1L). The 7-day acupuncture treatment that started after the completion of MPTP administration also exhibited partial but significant anti-parkinsonian effects (Fig. S1B-S1I). Taken together, acupuncture stimulation significantly alleviated both the PD-like motor and memory deficits in the MPTP mouse model.

**Fig. 1.**
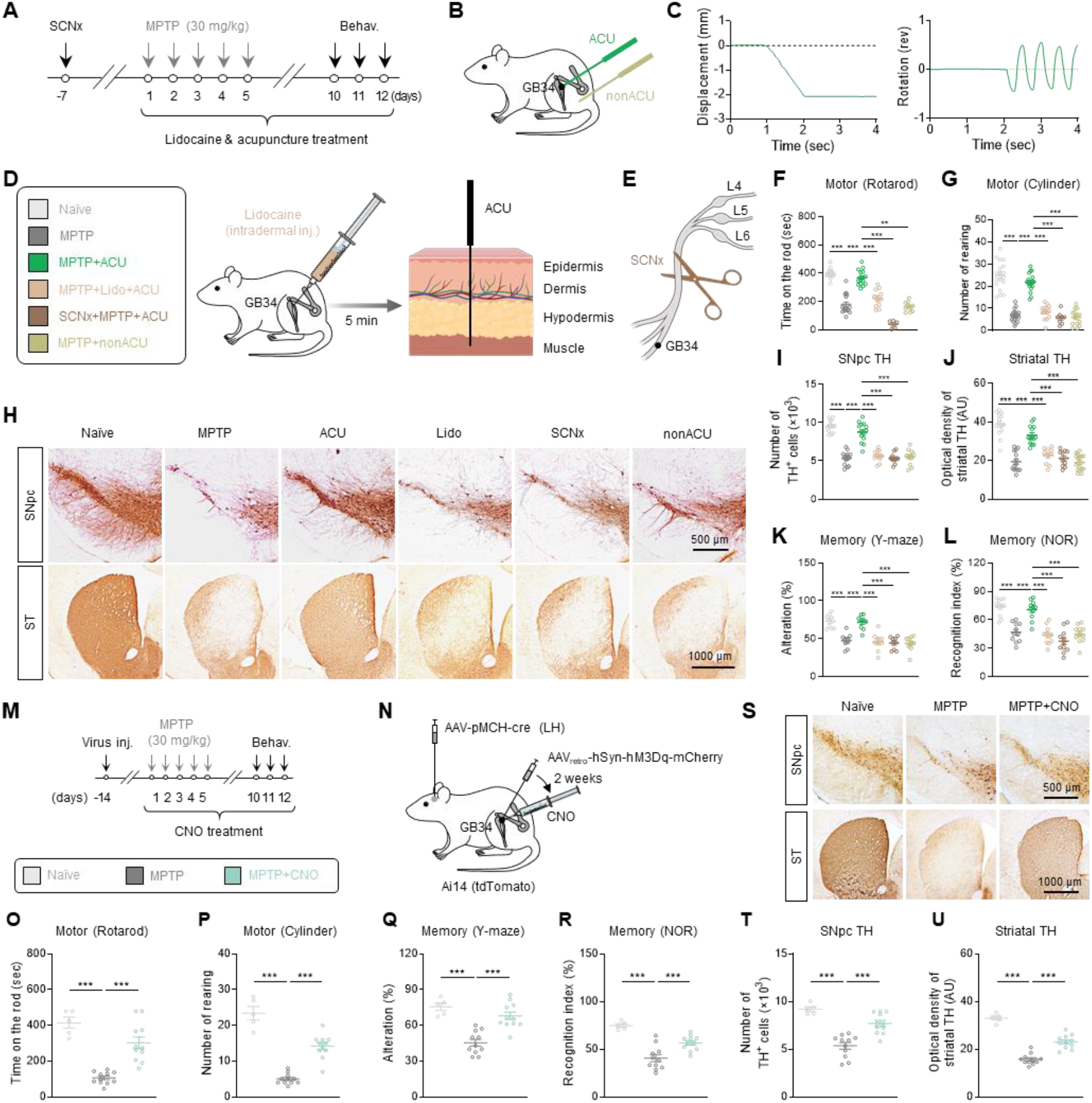
Acupuncture stimulation at GB34 elicits anti-parkinsonian effects through nerve conduction. **(A)** Experimental timeline of various interventions in the MPTP mouse model. **(B)** Schematic diagram of acupuncture treatment at GB34 acupoint and non-acupoint. **(C)** Quantitative measurement of displacement and rotation of the needle during acupuncture stimulation. **(D)** Left, group information. Right, schematic diagram of lidocaine-induced local nerve blockade 5 min prior to acupuncture stimulation. **(E)** Schematic diagram of sciatic nerve axotomy 7 days prior to acupuncture stimulation. **(F-G)** Motor function assessed by rotarod test (**F**) and cylinder test (**G**). Acupuncture stimulation alleviated the motor symptoms, which was blocked by lidocaine application and sciatic nerve axotomy. **(H)** Representative images of TH staining with SNpc and striatum tissues. **(I)** Numbers of TH-positive dopaminergic neurons in the SNpc. Quantification of optical density of striatal TH. Acupuncture stimulation alleviated the nigrostriatal TH loss, which was blocked by lidocaine application and sciatic nerve axotomy. Spatial working memory function assessed by Y-maze test. **(L)** Spatial memory function assessed by novel object recognition (NOR) test. Acupuncture stimulation alleviated the memory deficits, which was blocked by lidocaine application and sciatic nerve axotomy. **(M)** Experimental timeline of GB34 chemogenetic activation in the MPTP model and group information. **(N)** Schematic diagram of chemogenetic stimulation of peripheral afferent nerve fibers at acupoint GB34. **(O-P)** Motor function assessed by rotarod tests (**O**) and cylinder test (**P**) upon chemogenetic stimulation of acupoint GB34 in the MPTP model. **(Q-R)** Memory function assessed by Y-maze test (**Q**) and NOR test (**R**) upon chemogenetic stimulation of acupoint GB34 in the MPTP model. **(S)** Representative images of TH expression in the SNpc and striatum. **(T)** Numbers of TH-positive dopaminergic neurons in the SNpc. **(U)** Quantification of optical density of striatal TH. Statistical significance was assessed by Kruskal-Wallis ANOVA test with Dunn’s multiple comparison test **(F)** or one-way ANOVA with Tukey’s multiple comparison test. **p < 0.01. ***p < 0.001. All data are presented as mean ± SEM. Detailed statistical information is listed in **Table S1**.

Based on recent findings about the possible involvement of afferent nerve conduction in acupuncture effects (*24-26*), we also postulated that periphery-to-brain nerve conduction could mediate the anti-parkinsonian effects of acupuncture stimulation at GB34. To examine the necessity of this mechanism, we performed lidocaine-induced local nerve block (7 mg/kg; Fig. 1D) or axotomy of the sciatic nerve before acupuncture at GB34 (Fig. 1E). We found that both lidocaine application and sciatic nerve axotomy significantly blocked all observed anti-parkinsonian effects of acupuncture in the MPTP model (Fig. 1F-1L), suggesting that sensory nerve conduction from the peripheral acupoint is necessary for the effects of acupuncture stimulation at GB34. Next, we tested the sufficiency of afferent nerve conduction. We expressed hM3Dq, Gq-coupled designer receptors exclusively activated by designer drug (DREADD) (*27*), in the afferent nerve endings at GB34 using retro-adeno-associated virus (AAV) and activated it via intramuscular and subcutaneous injection of clozapine N-oxide (CNO) (1 mg/kg; Fig. 1M-1N). Intriguingly, we found that hM3Dq-mediated activation of afferent fibers located at GB34 significantly ameliorated PD-like motor symptoms, memory deficits, and nigrostriatal tyrosine hydroxylase (TH) loss in the MPTP model (Fig. 1O-1U). These findings indicate that activation of the afferent nerve endings at GB34 is necessary and sufficient to mimic the effects of acupuncture.

### Anatomical and functional link between acupoint and MCH^LH/ZI^ neurons

Based on previous studies demonstrating that the hypothalamic area, particularly the LH, is an important brain region that mediates the acupuncture effect (*14, 28-30*), we questioned whether acupoint GB34 is anatomically and functionally connected to the LH, especially MCH neurons. We first observed the significantly increased c-Fos expression in the MCH^LH/ZI^ neurons by chemogenetic activation of afferent fibers at the GB34 (Fig. S2A-S2D). Next, using pseudorabies virus (PRV)-green fluorescent protein (GFP), we first retrogradely traced neural circuits from GB34 to the brain in the pro-MCH (pMCH)-tdTomato mice, which were generated by viral expression of Cre under the pMCH-specific (>90%, Fig. S2A) promoter (AAV_dj_-pMCH-Cre) in the LH/ZI region of Ai14 reporter mice (Fig. 2A and S2E). We observed GFP^+^ neurons in the LH/ZI region, with a considerable proportion of them being tdTomato-expressing MCH neurons (Fig. 2B and 2C), which has previously been suggested as a possible contributor to the acupuncture effect (*14*). Bidirectional retrograde tracing from both GB34 and the LH/ZI region allowed us to further validate the neural connection from GB34 to the LH/ZI region. (Fig. S2F-S2H).

**Fig. 2.**
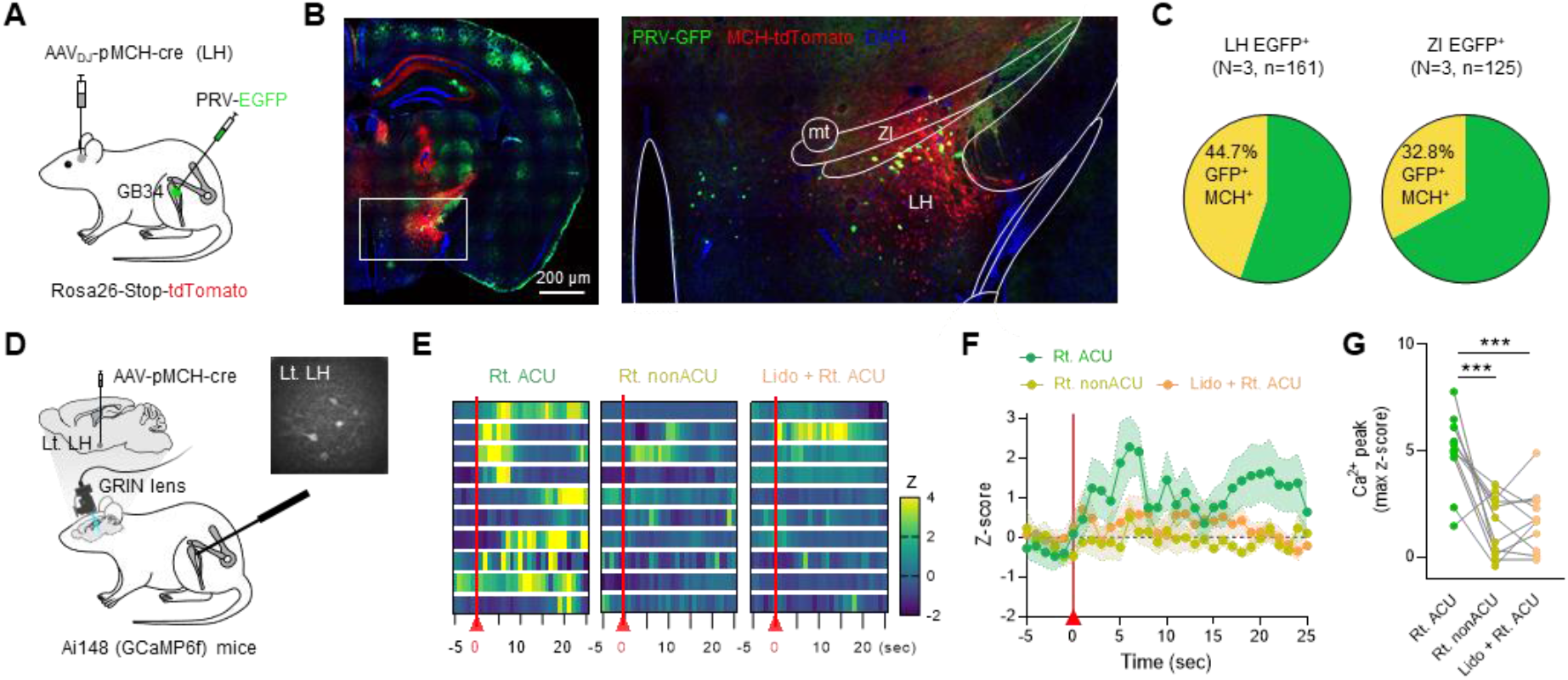
Anatomical and functional connection between a peripheral acupoint GB34 and MCH^LH/ZI^ neurons. **(A)** Schematic diagram of retrograde tracing of the neural path from acupoint to the brain using pseudorabies virus (PRV). **(B)** Identification of PRV-GFP-positive MCH neurons in the LH and ZI. **(C)** Quantification of PRV-GFP-positive MCH neurons. **(D)** Schematic diagram of *in-vivo* calcium imaging of MCH^LH/ZI^ neurons upon acupuncture stimulation. Right top, an example image of MCH-neuronal *GCaMP6f* signals. **(E)** Heatmaps of calcium signals from each neuron upon acupuncture or control stimulations. **(F)** Averaged time course of relative changes in *GCaMP6f* fluorescence indicating calcium signals. The average per group of the Z scored data is shown. **(G)** Quantification of peak amplitudes of calcium signals. Statistical significance was assessed by repeated-measure one-way ANOVA with Tukey’s multiple comparison test (**G**). **p < 0.01. ***p < 0.001. All data are presented as mean ± SEM. Detailed statistical information is listed in **Table S1**.

Next, we assessed the functional connectivity between GB34 and MCH^LH/ZI^ neurons by performing *in-vivo* Ca^2+^ imaging using single-photon microendoscopy upon acupuncture at GB34 in head-fixed and lightly anesthetized mice. We utilized Ai148 mice, which Cre-dependently express GCaMP6f (*31*), a genetically encoded calcium indicator (*32*), with AAV_dj_-pMCH-Cre injection into the LH/ZI regions (Fig. 2D). We found that acupuncture stimulation at GB34 elicited a significant Ca^2+^ rise in contralateral MCH^LH/ZI^ neurons, whereas needling at a non-acupoint or acupoint on the opposite hindlimb did not (Fig. 2E-2G and S2I-S2K). Together, these findings provided strong evidence of the anatomical and functional link between acupoint GB34 and MCH^LH/ZI^ neurons.

### MCH^LH/ZI^ neurons mediate anti-parkinsonian effects of acupuncture

To determine whether MCH^LH/ZI^ neurons are necessary for the anti-parkinsonian effects of acupuncture, we adopted a chemogenetic approach to selectively inhibit MCH^LH/ZI^ neurons using hM4Di, Gi-coupled DREADD (*27*) driven by the pMCH promoter, with systemic administration of CNO (1 mg/kg) (Fig. 3A and S3A). We found that the acupuncture-induced alleviation of motor symptoms, memory deficits, and nigrostriatal TH loss was fully blocked by hM4Di-mediated chemogenetic inhibition of MCH^LH/ZI^ neurons in the MPTP model (Fig. 3B-3H). The MPTP-induced spatial memory deficits were associated with impaired long-term potentiation (LTP) at the Shaffer Collateral-CA1 synapse of the HPC (Fig. 3I and 3J). And acupuncture treatment significantly reversed the LTP impairment, which was blocked by the chemogenetic inhibition of MCH^LH/ZI^ neurons. These findings indicate that MCH^LH/ZI^ neuronal activity is necessary for the therapeutic effects of acupuncture on both motor and non-motor symptoms.

**Fig. 3.**
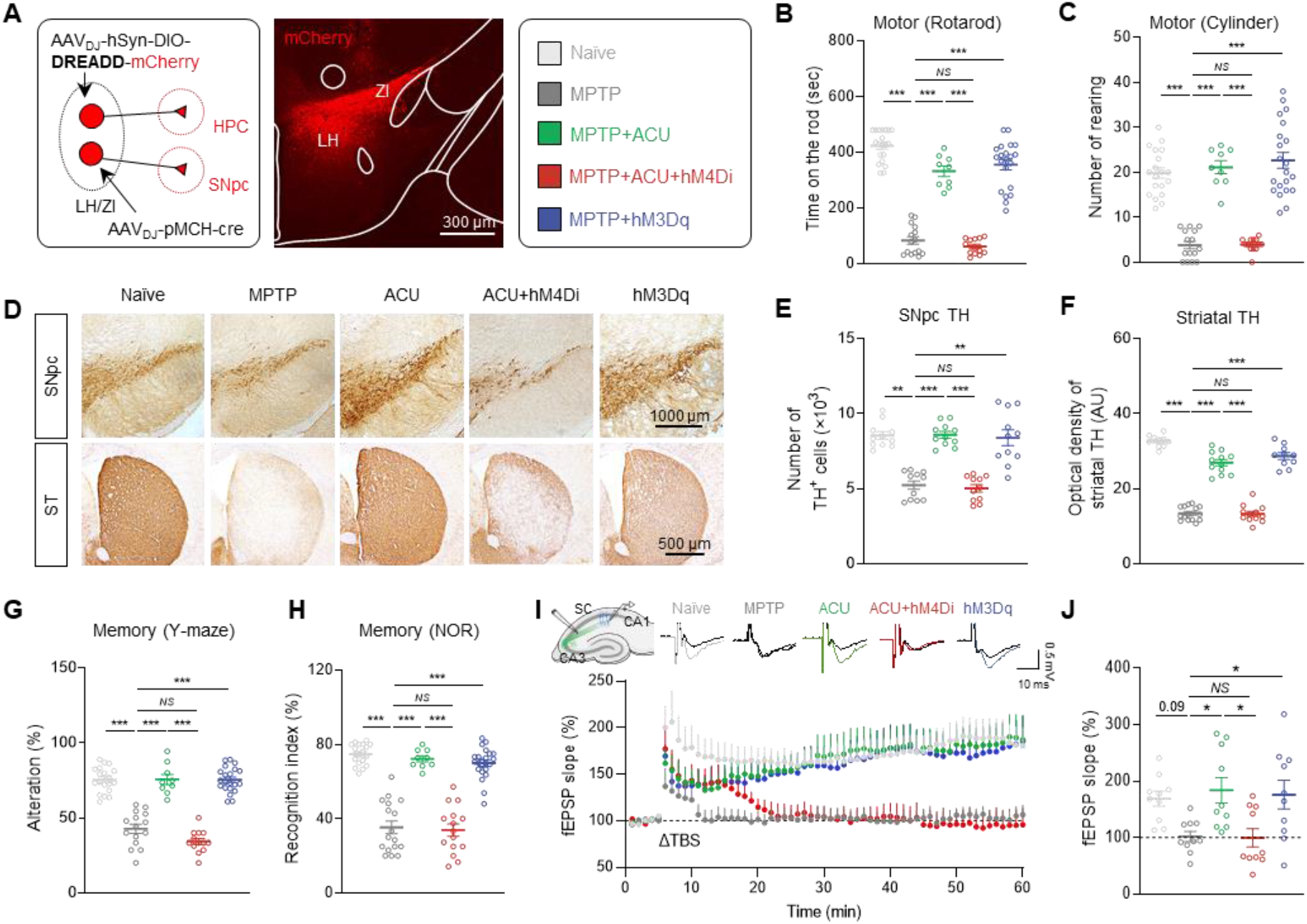
MCH^LH/ZI^ neuronal activity is critical for anti-parkinsonian effects of acupuncture. **(A)** Left, Schematic diagram of viral strategy for chemogenetic modulation of MCH^LH/ZI^ neurons. Middle, representative confocal image of *mCherry* expression in LH/ZI region. Right, group information. **(B-C)** Motor function assessed by rotarod tests (**B**) and cylinder test (**C**) upon chemogenetic manipulation of MCH^LH/ZI^ neurons with or without acupuncture treatment in the MPTP model. **(D)** Representative images of TH staining with SNpc and striatum tissues of each group. **(E)** Numbers of TH-positive dopaminergic neurons in the SNpc. **(F)** Quantification of optical density of striatal TH. **(G-H)** Memory function assessed by Y-maze test (**G**) and NOR test (**H**) upon chemogenetic manipulation of MCH^LH/ZI^ neurons with or without acupuncture treatment in the MPTP model. **(I)** Top, Representative fEPSP traces before (black) and 55 min after theta-burst stimulation (TBS). Bottom, time-course of fEPSP slope change. **(J)** Quantification of changes in the fEPSP slopes after TBS (for the last 10 min). Statistical significance was assessed by Kruskal-Wallis ANOVA test with Dunn’s multiple comparison test (**E**) or one-way ANOVA with Tukey’s multiple comparison test. **p < 0.01. ***p < 0.001. All data are presented as mean ± SEM. Detailed statistical information is listed in **Table S1**.

Next, we investigated whether MCH neurons are sufficient for the acupuncture effects by adopting hM3Dq-mediated chemogenetic activation (Fig. 3A and S3B). We found that the chemogenetic activation of MCH neurons without acupuncture recapitulated all observed aspects of the anti-parkinsonian effects of acupuncture in the MPTP model (Fig. 3B-3J), as well as in the A53T-mutated alpha-synuclein model (Fig. S3C-S3K) (*33*). MCH neuronal activation also alleviated MPTP-induced weight loss (Fig. S3L-S3N), consistent with previous findings (*34, 35*). Taken together, these findings indicate that MCH^LH/ZI^ neurons are necessary and sufficient for acupuncture-mediated alleviation of both PD-related motor and memory deficits.

### Two discrete subpopulations of MCH neurons: MCH^LH/ZI→SNpc^ and MCH^LH→HPC^

To investigate how MCH^LH/ZI^ neurons can mediate the effects of acupuncture on motor and memory deficits, we screened the projection of MCH^LH/ZI^ neurons using light-sheet imaging with pMCH-tdTomato mice (Fig. 4A, 4B, and S4A-S4B). We observed the tdTomato^+^ axon terminals in the GFP-labeled SNpc and HPC (Fig. 4B), two regions critically involved in motor and memory deficits in PD, respectively (*36-38*), in addition to other previously known target regions (Fig. S4C-S4E) (*39*). To date, it is unclear whether separate MCH^LH/ZI^ neurons project to distinct targets of the SNpc and HPC, or if a single MCH^LH/ZI^ neuronal population collateralizes onto multiple targets. To answer this question, we used an intersectional genetic approach to differentially label SNpc- or HPC-projecting MCH^LH/ZI^ neurons with mCherry and GFP, respectively (Fig. 4C). We found that mCherry^+^ SNpc-projecting MCH neurons were present throughout the LH and ZI (MCH^LH/ZI→SNpc^), while GFP^+^ HPC-projecting MCH neurons were clustered within the LH (MCH^LH→HPC^) (Fig. 4D and 4E). Moreover, GFP^+^ and mCherry^+^ neurons were mostly separated (Fig. 4E). We also observed no significant GFP^+^ fibers in the SNpc or mCherry^+^ fibers in the HPC (Fig. 4D). These findings indicate that MCH^LH/ZI→SNpc^ and MCH^LH→HPC^ neurons are anatomically distinct neuronal populations that represent separate neural pathways.

**Fig. 4.**
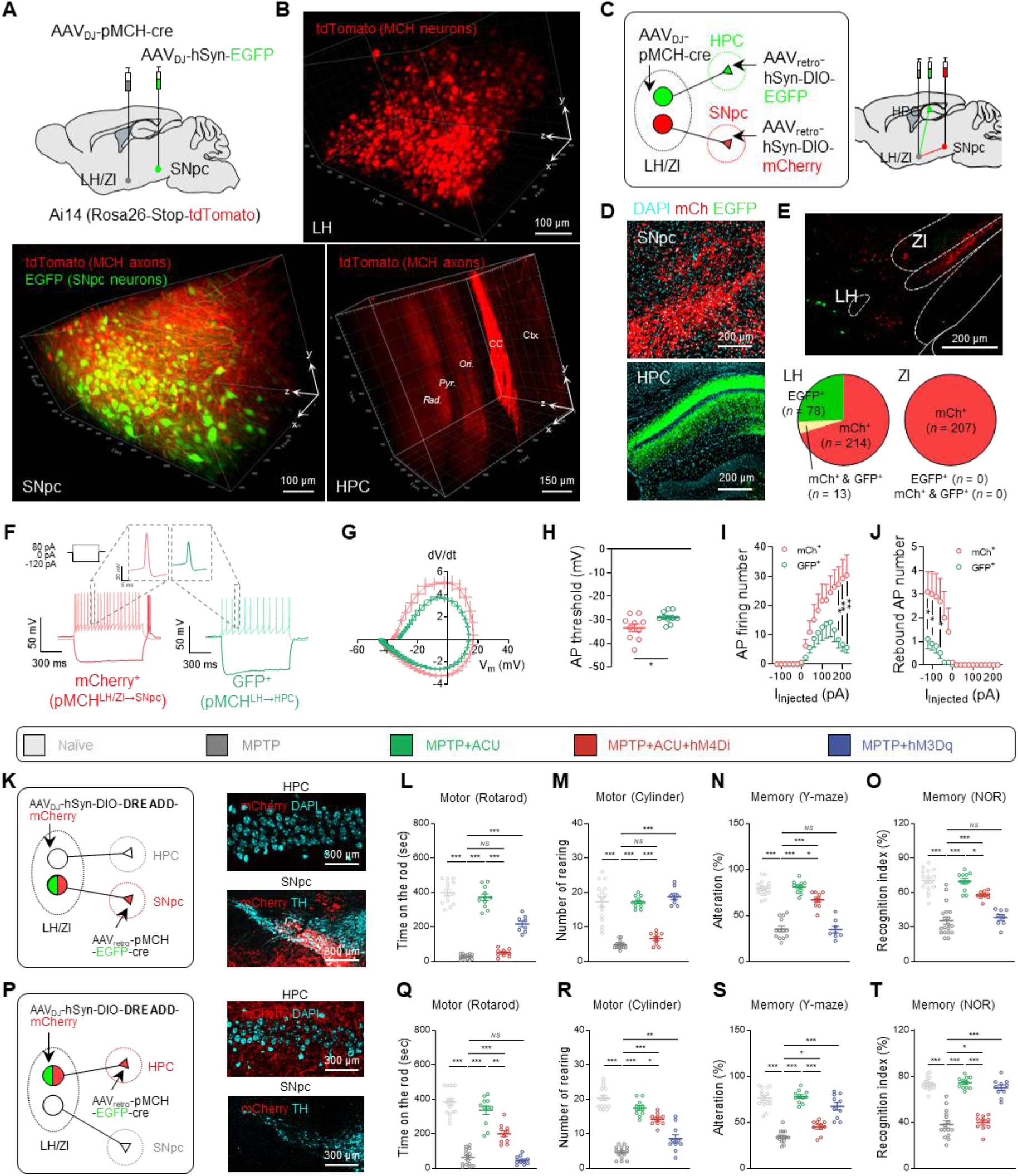
Discrete MCH projections, MCH^LH/ZI→SNpc^ and MCH^LH→HPC^, are responsible for motor and memory function, respectively. **(A)**Schematic diagram of viral injections for elucidating the MCH^LH/ZI^ neuronal projections. Three-dimensional rendering of a cleared mouse brain showing *tdTomato*-labeled MCH^LH/ZI^ neuronal soma (top) and projections to SNpc and HPC (bottom). **(C)** Schematic diagram of viral strategy for differentially labeling MCH neurons projecting to SNpc (*mCherry*) or HPC (EGFP). **(D)** Representative confocal images of MCH axon fibers in SNpc or HPC originated from the soma located in LH/ZI. **(E)** Intra-LH/ZI localization and quantification of MCH neuronal subpopulations projecting to SNpc (*mCherry*; MCH^LH/ZI→SNpc^) or HPC (EGFP; MCH^LH→HPC^). **(F)** Representative traces of membrane potentials recorded from MCH^LH/ZI→SNpc^ or MCH^LH→HPC^ neurons upon current steps (left top). **(G)** Phase plot analyses of action potentials. **(H-J)** Intrinsic electrophysiological properties of MCH^LH/ZI→SNpc^ or MCH^LH→HPC^ neurons; AP threshold (**H**), AP firing numbers upon depolarizing current step (**I**), rebound AP numbers upon hyperpolarizing current steps (**J**). **(K, P)** Schematic of viral strategy for projection-specific chemogenetic manipulation of MCH^LH/ZI→SNpc^ (**K**) or MCH^LH→HPC^ (**P**). **(L, M, Q, R)** Motor function assessed by rotarod tests (**L, Q**) and cylinder test (**M, R**). **(N, O, S, T)** Memory function assessed by Y-maze test (**N, S**) and NOR test (**O, T**). Statistical significance was assessed by unpaired two-tailed Student’s t-test (**H-J**) or one-way ANOVA with Tukey’s multiple comparison test. *p < 0.05, ** p < 0.01, ***p < 0.001, NS, non-significant. All data are presented as mean ± SEM. Detailed statistical information is listed in **Table S1**.

Next, we examined the electrophysiological properties of these two subpopulations by performing *ex-vivo* whole-cell patch-clamp. We found that the distinct action potential (AP) firing patterns (Fig. 4F-4J, S4F-S4J). MCH^LH→HPC^ neurons were characterized by lower AP firing probability (Fig. 4H and 4I), while MCH^LH/ZI→SNpc^ neurons exhibited typical rebound low-threshold bursting after hyperpolarization (Fig. 4J). Taken together, our findings provided strong evidence of two discrete subpopulations of MCH^LH/ZI^ neurons: MCH^LH/ZI→SNpc^ and MCH^LH→HPC^.

To investigate whether these projections form functional circuits with target neurons in the SNpc and HPC, we performed *ex-vivo* Ca^2+^ imaging of SNpc dopaminergic neurons and CA1 pyramidal neurons upon chemogenetic activation of MCH^LH/ZI^ neurons. We found that bath application of CNO (5 μM) was sufficient to induce a significant calcium increase in SNpc dopaminergic neurons (Fig. S5A-S5D) and CA1 pyramidal neurons (Fig. S5E-S5H). These results together indicate that distinct subpopulations of MCH^LH/ZI^ neurons send functional projections to SNpc and CA1.

### MCH^LH/ZI→SNpc^ and MCH^LH→HPC^ differentially mediate motor and memory function

Since we demonstrated that MCH neurons distinctly project to the SNpc or HPC, which are the crucial brain regions of PD-related motor and memory deficits, respectively, we raised the possibility that the acupuncture effects can be mechanistically dissected at the circuit level. Thus, to determine whether two anatomically discrete MCH neuronal projections differentially mediate the effects on motor and memory deficits, we expressed hM4Di or hM3Dq exclusively in MCH^LH/ZI→SNpc^ or MCH^LH→HPC^ neurons in a cell type- and projection-specific manner, by adopting a Cre-dependent retrograde labeling strategy (Fig. 4K and 4P). We found that the chemogenetic inhibition of MCH^LH/ZI→SNpc^ neurons significantly blocked the effects of acupuncture on motor dysfunction and TH loss, whereas the effects of acupuncture on memory deficits was minimally affected. At the same time, chemogenetic activation of MCH^LH/ZI→SNpc^ neurons significantly mimicked the effects of acupuncture on motor dysfunction and TH loss, while memory deficits were not alleviated (Fig. 4K-4O and S6A-S6D). In contrast, the chemogenetic inhibition of MCH^LH→HPC^ neurons fully abolished the effects of acupuncture on memory function, but much less on motor function. The chemogenetic activation of MCH^LH→HPC^ neurons significantly recapitulated the effects of acupuncture on memory function, but not on motor function and the associated nigrostriatal TH expression (Fig. 4P-4T and S6E-S6H). Collectively, these results indicate that the two discrete MCH neuronal projections to the SNpc or HPC are differentially responsible for motor and memory recovery in the MPTP model.

### MCH and MCHR1 are critical for the acupuncture effect

Since MCH neuronal activity is critical for the anti-parkinsonian effects of acupuncture, we tested whether the neuropeptide MCH and its receptor MCHR1 are responsible for the effects. We first pharmacologically blocked MCHR1 by intraperitoneal administration of a selective and brain-penetrable MCHR1 antagonist, TC-MCH7c (10 mg/kg). We found that the acupuncture-mediated alleviation of motor and memory deficits, as well as nigrostriatal TH loss, was not observed in the MPTP model when MCHR1 was blocked (Fig. 5A-5E and S7A-S7C). Consistently, intranasal administration of MCH (0.5 μg in 30 μL saline) significantly recapitulated the acupuncture effects on MPTP-induced motor deficits, TH loss, and memory impairments (Fig. 5A-5E and S7A-S7C). We also tested whether the region-specific gene-silencing of MCHR1 by short-hairpin RNA (shRNA) (Fig. S7D-S7I) interfered with the anti-parkinsonian effects of acupuncture (Fig. 5F-5J). Intriguingly, we found that MCHR1 gene-silencing in the SNpc blocked the acupuncture-mediated improvement of motor function but did not affect memory function in the MPTP model. Likewise, MCHR1 gene-silencing in the HPC hindered acupuncture-mediated memory improvement, while the acupuncture effects on motor function and nigrostriatal TH expression were less affected (Fig. 5G-5J and S7J-S7N). These findings together indicate that MCHR1-mediated MCH signaling is indeed critical for the acupuncture effect.

**Fig. 5.**
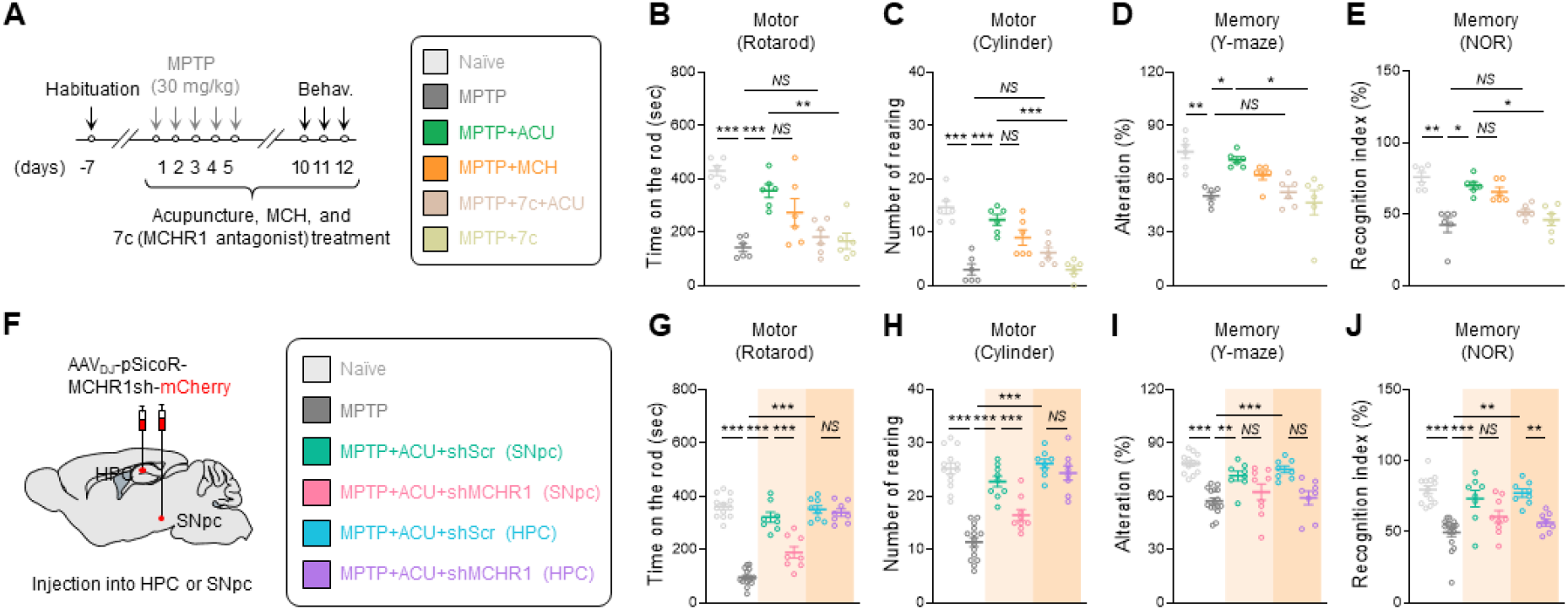
MCH-MCHR1 activation is critical for multi-therapeutic effects of acupuncture. **(A)** Left, experimental timeline of various interventions including acupuncture, intranasal administration of MCH, and intraperitoneal administration of MCHR1 antagonist in the MPTP model. Right, group information. **(B, C)** Motor function assessed by rotarod tests (**B**) and cylinder test (**C**) upon various interventions in the MPTP model. **(D, E)** Memory function assessed by Y-maze test (**D**) and NOR test (**E**) upon various interventions in the MPTP model. **(F)** Schematic diagram of AAV-mediated gene-silencing of MCHR1. **(G, H)** Motor function assessed by rotarod tests (**G**) and cylinder test (**H**). **(I, J)** Memory function assessed by Y-maze test (**I**) and NOR test (**J**). Statistical significance was assessed by Kruskal-Wallis ANOVA test with Dunn’s multiple comparison test (**I, J**) or one-way ANOVA with Tukey’s multiple comparison test. *p < 0.05, ** p < 0.01, ***p < 0.001, NS, non-significant. All data are presented as mean ± SEM. Detailed statistical information is listed in **Table S1**.

### Acupuncture and MCH neuronal activation commonly alter transcriptomic profiling

Next, we performed RNA-sequencing to unbiasedly explore the common mode-of-action of acupuncture and chemogenetic activation of MCH neurons in alleviating the PD-related motor and memory deficits. To investigate alterations in gene expression profiles across the four groups (naïve, MPTP, MPTP+Acupuncture, and MPTP+hM3Dq), we used substantia nigra (SN) and HPC tissues from each mouse (N = 3 for each group; Fig. S8A and S10A). We analyzed the RNA-sequencing data using fragments per kilobase of transcript per million mapped reads (FPKM; >1; Fig. S8B-S8H). We found that the MPTP model showed reduced expression of dopaminergic neuron genes (Fig. 6A and S8F) and increased expression of several marker genes of reactive astrocytes (Fig. 6B and S8G) and microglia (Fig. 6C and S8H), which were generally reversed by both acupuncture and MCH neuronal activation (Fig. 6A-6C and S8C-S8H). We further determined differentially expressed genes (DEGs) with a cutoff of an adjusted p-value of less than 0.05 and a fold-change of greater than 1.5 between the two groups. We identified 171 SNpc DEGs that were increased by MPTP treatment and reversed by both acupuncture and hM3Dq-activation of MCH neurons (MPTP-up, ACU-down, and hM3Dq-down) and 81 DEGs that were changed in the opposite direction (MPTP-down, ACU-up, and hM3Dq-up). Notably, gene ontology (GO) analysis of these SN DEGs supported the common therapeutic effects of acupuncture and MCH neuronal activation the in MPTP model (Fig. S9).

**Figure 6.**
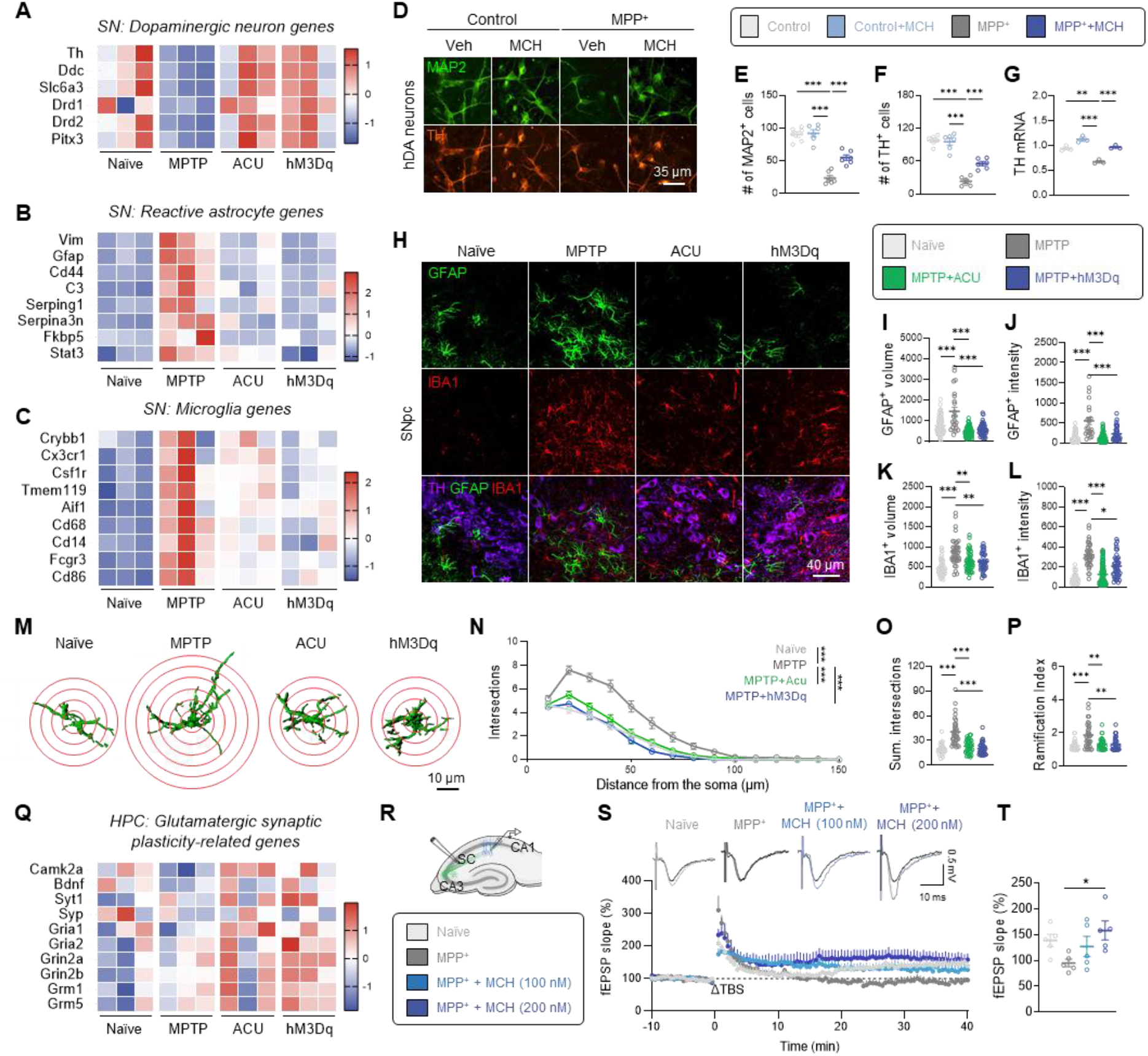
Acupuncture and MCH^LH/ZI^ activation share transcriptomic signatures. **(A-C)** Heatmap showing enriched genes in specific cell types such as DA neurons (**A**), reactive astrocytes (**B**), and microglia (**C**) in the SN. **(D)** Immunostaining with MAP2 and TH in hDA neurons treated with MPP^+^ and MCH. **(E, F)** The number of MAP2-positive (**E**) and TH-positive cells (**F**). **(G)** mRNA expressions of *TH*. **(H)** Representative confocal images of GFAP-positive astrocyte and IBA1-positive microglia in the SNpc. **(I, J)** Quantification of volume (**I**) and GFAP intensity (**J**) of each astrocyte. **(K, L)** Quantification of volume (**K**) and IBA1 intensity (**L**) of each microglia. **(M)** Three-dimensionally rendered images of GFAP-positive astrocytes. Red circles indicate superimposed spheres centered around astrocyte somata used for Sholl analysis. **(N-P)** Morphometric analyses of GFAP-positive astrocytes by Sholl analysis; Sholl intersections by distance from the soma (**N**), total number of intersections (**O**), and ramification index (**P**). **(Q)** Heatmap showing glutamatergic synaptic plasticity-related genes in the HPC. **(R)** Schematic diagram of *ex-vivo* field recording for LTP measurement at SC-CA1 synapse. **(S)** Top, Representative fEPSP traces before (black) and 40 min after theta-burst stimulation (TBS). Bottom, time-course of fEPSP slope change. **(T)** Quantification of changes in the fEPSP slopes after TBS (for the last 10 min). Statistical significance was assessed by Kruskal-Wallis ANOVA test with Dunn’s multiple comparison test (**K, L, O, P**), two-way ANOVA with Tukey’s multiple comparison test (**N**), or one-way ANOVA with Tukey’s multiple comparison test. *p < 0.05, ** p < 0.01, ***p < 0.001, NS, non-significant. All data are presented as mean ± SEM. Detailed statistical information is listed in **Table S1**.

The increase in dopaminergic neuron-related genes by acupuncture and MCH neuronal activation was recapitulated by direct MCH application onto human induced pluripotent stem cell (hiPSC)-derived dopaminergic (hDA) neurons which exhibits PD-like phenotypes by 1-Methyl-4-phenylpyridinium (MPP^+^) treatment (Fig. 6D-6G and S9E-S9G). This result suggests that MCH can elicit neuroprotective action in a cell-autonomous manner. On the other hand, the reduction in gliosis-related transcriptomic profiling in the SN tissue, particularly astrogliosis, is also noteworthy because previous reports have demonstrated that reactive astrocytes can lead to dopaminergic neurodegeneration in a non-cell-autonomous manner in PD pathology (*33, 40-42*). Our immunohistochemistry results demonstrated that the intensities of GFAP and IBA1, as well as cellular volumes of astrocytes and microglia, were significantly increased in the SNpc of the MPTP model, which were reversed by both acupuncture and chemogenetic activation of MCH neurons (Fig. 6H-6L). Additionally, the high ramification of astrocytes, a key morphological characteristic of reactive astrocytes, was significantly reduced by both acupuncture and chemogenetic activation of MCH neurons (Fig. 6M-6P). These findings indicate that acupuncture and the associated MCH neuronal activation can alleviate the MPTP-induced reactive gliosis. Taken together, these findings suggest that MCH can restore the dopaminergic neurons via both non-cell-autonomous and cell-autonomous mechanisms, which is responsible for the anti-parkinsonian effect of acupuncture.

In the HPC, we observed a reduction in the expression of genes related to glutamatergic synaptic plasticity and homeostatic microglial action, whereas there was an increasing trend in the expression of genes related to reactive astrocytes and pro-inflammatory microglia following MPTP treatment. Acupuncture and hM3Dq-mediated MCH neuronal activation appeared to reverse these changes (Fig. 6Q and S10). Particularly, GO analysis with 27 HPC DEGs (MPTP-up, ACU-down, and hM3Dq-down) demonstrated that inflammatory signaling was attenuated by acupuncture and MCH neuronal activation (Fig. S11). These findings indicate that acupuncture and chemogenetic activation of MCH neurons commonly alters the transcriptomic profiling of the SN and HPC of the MPTP model towards the normal state. Interestingly, we revealed that MCH was sufficient to recapitulate the effects of acupuncture and hM3Dq-mediated MCH neuronal activation in restoring the hippocampal synaptic plasticity (Fig. 6R-6T). These findings strongly suggest that MCH is the essential molecular substance for the anti-parkinsonian effects of acupuncture in the brain. Altogether, our findings strongly suggest that the anti-parkinsonian effects of acupuncture are attributed to MCH-mediated alleviation of reactive gliosis, protection of dopaminergic neurons in the SNpc, and enhancement of glutamatergic synaptic plasticity in the hippocampus.

## Discussion

The neurobiological mechanisms underlying how acupuncture stimulation at a peripheral acupoint modulates neural activity in the brain has remained unclear for several millennia. Our study demonstrated that acupuncture stimulation at a hindlimb acupoint GB34 activates MCH^LH/ZI^ neurons via afferent nerve conduction. Through cell type- and projection-specific chemogenetic manipulation with an intersectional genetic approach, we demonstrated that acupuncture can simultaneously alleviate the PD-related motor and memory deficits through MCH neuronal activation. We further identified two subpopulations of MCH^LH/ZI^ neurons and delineated their discrete functional relevance: motor behaviors (MCH^LH/ZI→SNpc^) and non-motor behaviors (MCH^LH→HPC^). This finding is noteworthy because it has not been studied whether the separate subpopulations of MCH^LH/ZI^ neurons have distinct projections involving differential functions in the system level, despite recent highlights on the diversity of several hypothalamic neurons (*43-45*). Our findings strongly suggest that these two distinct circuits from MCH^LH/ZI^ neurons are the cellular and circuit-level neuroanatomical basis of the multi-function of acupuncture in PD (Fig. S12).

We have revealed that the anti-parkinsonian effects of acupuncture in both PD-related motor and non-motor symptoms were attributed to the action of MCH. First, while the motor symptoms of PD are known to result from the extensive nigrostriatal dopaminergic neuronal death, the non-cell-autonomous mechanisms through reactive astrocytes and microglia have also been reported (*33, 41, 46*). Our study demonstrates that MCH can protect dopaminergic neurons both cell-autonomously and non-cell-autonomously via reducing reactive gliosis. Second, while the cardinal motor symptoms have been emphasized as the primary target for most existing PD treatments, the non-motor symptoms of PD, including cognitive impairment, have received little attention, despite their clinical importance. This study demonstrates that acupuncture treatment significantly recovers the long-term potentiation at the Schaffer collateral-CA1 synapse at the hippocampus and the spatial memory impairment in PD mice, through the MCH-MCHR1 interaction. This is in line with previous findings showing that MCHR1 mRNA is highly expressed in the CA1 hippocampus (*47, 48*) and that MCH might contribute to hippocampal synaptic plasticity and memory behaviors through the signaling pathway involving brain-derived neurotrophic factor (BDNF) and its receptor, tropomyosin receptor kinase B (TrkB) (*49-51*). Taken together, our results propose MCH as the potential therapeutic target as well as molecular basis of anti-parkinsonian effects of acupuncture on both motor and non-motor symptoms.

Current PD therapies have only focused on symptomatic relief through dopamine supplementation or dopamine degradation blockade, which cannot modify the disease progression. However, our findings suggest that MCH may have the potential to modify the disease progression of PD through its ability to promote dopaminergic neuroprotection and reduce reactive gliosis. We further propose peripheral nerve stimulation by acupuncture could be an effective strategy for eliciting MCH^LH/ZI^ neuronal activity to relieve motor and non-motor symptoms in PD. Taken together, our study proposes the neuroanatomical basis of the anti-parkinsonian effects of acupuncture, which was previously considered a non-scientific intervention. These findings shed light on an exciting new avenue for PD therapy.

## Acknowledgements

Graphical abstract was created via Biorender.com.

## Author contributions

Conceptualization: MHN, HJP

Methodology: HYK, SEL, SEL, SJO, JGK, CJL

Investigation: JYO, HL, SYJ, HJK, GP, AS, JHJ, JK (Junyeop Kim), SKY, MSS, SEL, YEH, TYH

Visualization: JYO, HL, MHN

Resources: SEL, JK (Jeongjin Kim), JK (Jongpil Kim), SJO, CJL, MHN, HJP Funding acquisition: MHN, HJP

Project administration: MHN, HJP

Supervision: CJL, MHN, HJP

Writing - Original Draft: JYO, HL, MHN

Writing - Review & Editing: JYO, HL, MHN, HJP

## Data and materials availability

The data supporting the findings from this study are available within the manuscript and its Supplementary Information. The RNA-sequencing data has been deposited in the National Center for Biotechnology Information (NCBI) Sequence Read Archive (SRA) database with the BioProject accession code PRJNA909842. Because the data is not publicized until publication of the current manuscript, we provide the access link for the reviewers (https://dataview.ncbi.nlm.nih.gov/object/PRJNA909842?reviewer=6jslojpfaf9lvp5aoa1sog7ihg). Source data for every graph are provided with this paper.

## Supplementary Materials

Materials and methods

Figs. S1 to S12

Tables S1 to S3

References (52_–_61)

