## Supplementary material for "Discrete projections from MCH neurons mediate anti-parkinsonian effects of acupuncture": 230604 supple data STM V3 nam - biorxiv.pdf

**The PDF file includes:**

Materials and Methods
Figs. S1 to S12
Tables S1 to S3
References and Notes (52 – 61)

### Materials and Methods

Mice

All animal experimental procedures were conducted in accordance with the National Institutes of Health guidelines and approved by the Institutional Animal Ethical Committee, Kyung Hee University (Seoul, Korea, Approval Number KHSASP-19-412), and Korea Institute of Science and Technology (KIST; Seoul, Korea, Approval Number KIST-2020-177). If there is no specific explanation, 8–10-week-old male *C57BL/6J* mice were used for experiments. *Ai148* (B6.Cg-$Igs7^{tm148.1(tetO\text{-}GCaMP6f,CAG\text{-}tTA2)Hze}$/J) transgenic mice were used for *in-vivo* and *ex-vivo* $Ca^{2+}$ imaging. *CamK2a-Cre* transgenic mice crossed with *Ai148* transgenic mice were used for *ex-vivo* $Ca^{2+}$ imaging of pyramidal neurons in the CA1 HPC. *Ai14* (B6;129S6-$Gt(ROSA)^{26Sortm14(CAG\text{-}tdTomato)Hze/}$J) transgenic reporter mice were used for visualizing the MCH neurons and their projections with $AAV_{DJ}$*-pMCH-cre* virus injection. All mice were co-housed at the animal facility with a temperature of 21 ± 2 ºC, a relative humidity of 50 ± 15%, and a 12 h light/dark cycle (lights on and off at 8 am and 8 pm). All mice were provided with food and water *ad libitum.* All mice were randomly allocated to each experimental group. All experiments were performed with age-matched controls and different sets of animals were used for each experiment.

Virus preparation and stereotaxic injection

For all surgical procedures, mice were anesthetized with 2% isoflurane and head-fixed in a mouse stereotaxic apparatus (#68513, RWD, USA; Model 940, KOPF, USA). All virus injections were delivered at a rate of 0.2 μL $min^{-1}$ via a blunt-end injection needle (33-gauge, CMA, USA) using a microinjection syringe pump (KDS 310, KD Scientific, USA).

To visualize the MCH neurons or chemogenetically manipulate their activity, we injected $AAV_{DJ}$*-pMCH-EGFP-cre*, $AAV_{DJ}$*-pMCH-cre*, $AAV_{DJ}$*-hSyn-DIO-hM3Dq-mCherry*, or $AAV_{DJ}$*-hSyn-DIO-hM4Di-mCherry* into the LH (AP = -1.0 mm, ML = ±1.2 mm from bregma, DV = -4.85 mm from dura). The viruses injected into the LH were repeatedly spread to the ZI. The viruses injected into the LH were spread to the ZI. To visualize the $MCH^{LH\rightarrow HPC}$ or $MCH^{LH/ZI\rightarrow SNpc}$ projection or retrogradely express *Cre* in $MCH^{LH\rightarrow HPC}$ or $MCH^{LH/ZI\rightarrow SNpc}$ neurons, we injected $AAV_{retro}$*-hSyn-DIO-EGFP*, $AAV_{retro}$*-hSyn-DIO-mCherry* or $AAV_{retro}$*-pMCH-EGFP-cre* into the CA1 HPC (AP = -1.94 mm, ML = ±1.5 mm from bregma, DV = -1.3 mm from dura) or SNpc (AP = -3.2 mm, ML = ±1.25 mm from bregma, DV = -4.0 mm from dura). For *ex-vivo* $Ca^{2+}$ imaging of dopaminergic neurons in the SNpc, $AAV_{DJ}$*-DDC-cre* was injected into the SNpc of *Ai148* transgenic mice. To generate the viral *A53T* overexpression-induced PD mouse model (*33*), we injected $AAV_{DJ}$*-CMV-A53T-SNCA* into the SNpc. For gene-silencing of MCHR1 expression in HPC and SNpc, we injected *AAVDJ-pSicoR-MCHR1sh-mCherry* into CA1 HPC or SNpc. All AAVs were manufactured by the KIST Research Animal and Resource Center (KIST RARC). For viral trans-synaptic tracing from GB34 to the DRG, spinal cord, and brain, PRV-CMV-EGFP (P01001, BrainVTA, China) and PRV-CMV-RFP (P01002, BrainVTA) were utilized. We injected 1.5 uL of PRV-CMV-EGFP into the LH/ZI region and PRV-CMV-RFP was injected into GB34. PRV-CMV-GFP was injected into the GB34 with three different depths: 2 mm (muscular layer), 1.5 mm, and 1 mm (subcutaneous layer). All PRVs were obtained from BrainVTA (http://www.brainvta.tech). The detailed information about virus titers, volumes, and total amount for each experiment is listed in Table S2.

MPTP-induced parkinsonian model in mice

All MPTP experiments used the subchronic regimen consisting of a daily intraperitoneal (i.p.) injection of MPTP (30 mg $kg^{-1}$ per day, 23007-85-4, Sigma-Aldrich, USA) for five consecutive days. Control animals received saline injections only.

Acupuncture treatment

Acupuncture treatment was performed 2 h after MPTP injection for 12 consecutive days by inserting as stainless-steel acupuncture needle (15 mm in length, 0.20 mm diameter, Haeng-lim-seo-weon Acuneedle Co, Korea) with a depth of about 3 mm bilaterally at the GB34 acupoint. The GB34 acupoint, located at the intersection point of lines from the anterior borders to the head of the fibula, has been used to treat movement disorders in traditional East Asian medicine, and recent neuroimaging studies have confirmed its effect on motor function (*6, 58, 59*). Acupuncture stimulation at GB34 acupoint has been reported to be effective for alleviating motor symptoms in PD animal models (*13, 23, 60*). We chose a control non-acupoint (nonACU) located at the point about 3 mm lateral side of a tail on the gluteus muscle. These inserted needles were then turned at a rate of two spins per sec for 15 sec and removed immediately. Treatment was performed accurately and quickly to minimize stress on the mice. Also, mice in all groups were mildly immobilized by holding their necks, with the head in an upright position for 30 sec to give the same immobilization stress as the acupuncture group.

Sciatic axotomy and lidocaine treatment

For sciatic nerve transection, mice were anesthetized with rompun (100 μL, i.p.; Bayer, Korea) and 2% zoletil (150 μL, i.p.; Virbac S.A, France). The bilateral hind thigh was shaved and received a skin incision at mid-thigh level and the sciatic nerve was exposed. Then, the nerves were transected, and a ligature was performed in the proximal segment in order to prevent spontaneous reinnervation. For blocking the axonal reflex with a local anesthetic, 10 μL lidocaine (lidocaine-HCl 400 mg; Huons, Korea) was intradermally injected at the acupoint 5 min before acupuncture stimulation.

Chemogenetic manipulation

To manipulate the activity of MCH neurons in LH/ZI, we adopted *Cre*-dependent Designer Receptors Activated Only by Designer Drug (DREADD) expression and i.p. administration of CNO. Specifically, we injected $AAV_{DJ}$*-pMCH-cre* virus into LH/ZI to express *Cre* specifically in MCH neurons. Simultaneously, we injected $AAV_{DJ}$*-hSyn-DIO-hM4Di-mCherry* or $AAV_{DJ}$*-hSyn-DIO-hM3Dq-mCherry* virus into LH/ZI to *Cre*-dependently express Gi-DREADD or Gq-DREADD in the *Cre*-induced $MCH^{LH/ZI}$ neuron, respectively. Two weeks later, we started to administer MPTP. To activate Gi-DREADD and Gq-DREADD, we intraperitoneally administered CNO at a dose of 1 mg $kg^{-1}$ for 12 days. In terms of Gi-DREADD-mediated inhibition, we treated with CNO 30 min before acupuncture stimulation. For circuit-specific manipulation, we injected $AAV_{retro}$*-hSyn-DIO-hM4Di-mCherry* and $AAV_{retro}$*-hSyn-DIO-hM3Dq-mCherry* virus into HPC or SNpc.

For chemogenetic activation of sensory afferents at GB34 acupoint, we injected *$AAV_{retro}$-hSyn-hM3Dq-mCherry* into bilateral GB34 at the three different depths of 1, 1.5, and 2 mm for targeting subcutaneous, superficial, and deep muscular layers to mimic the acupuncture effects by chemogenetic activation of afferent nerve fibers surrounding the GB34 acupoint. Two weeks later, we injected CNO (1 mg/kg, 50 uL in total) into the bilateral GB34.

Administration of MCH and MCHR1 blocker

To assess the possible therapeutic effects of MCH in the MPTP mouse model, mice were intranasally administered with MCH (0.5 μg in 30 μL saline; #3806, Tocris, UK) for 12 consecutive days 2 h after MPTP treatment. To pharmacologically inhibit MCHR1, 10 mg of MCHR1 antagonist TC-MCH7c (7c; 10 mg/kg, dissolved in 8 mL of 1% dimethyl sulfoxide) was intraperitoneally administered to the mice 30 min before the acupuncture treatment.

Motor behavioral assays

*Rotarod test.* An accelerated rotarod test was conducted to evaluate the coordination and balance of motor function (*61*). On the day of testing, mice were kept in their home cages and acclimated to the testing room for at least 30 min. A 2-min acclimation session at 2 rpm was performed on the first day before the test phase. Rotarod testing (MED Associates, Inc, USA) involves placing mice on a rotating bar and determining the length of time that they can retain their balance while the rotation speed increases over 480 s. The speed is increased from 3.5 to 35 rpm, reaching maximum speed at 5 min. Each trial was terminated when the mouse fell off. The latency to fall from the rotating rod was scored by automatic timers and falling sensors on the rotarod.

*Cylinder test.* A cylinder test was performed to evaluate the spontaneous explorative behavior in a new environment (*62*). Mice were placed in a transparent plastic cylinder (12 cm in diameter × 20 cm tall) for 1 min before the experiment. After adaptation, when mice try to explore different areas of the cylinder by standing on their hindlimbs and leaning with the forelimbs on the cylinder wall, the cylinder wall touches (numbers) were counted by observers for 3 min.

*Adhesive removal test.* The adhesive removal test was established in α-synuclein *A53T* mice to determine the effects of MCH activation on sensory-motor behavior (*63*). Mice were placed in a clean cage and allowed to habituate for 30 min. A small circular adhesive paper sticker (5 mm in diameter) was gently but firmly attached to the snout. The ability to remove the adhesive sticker was evaluated by the time required for mice to remove the sticker. Mice were given 60 s to complete this sensorimotor task. Each mouse was subjected to three trials separated by 1 min rest and the results were averaged.

Memory behavioral assays

*Y-maze test.* Spatial working memory was measured by spontaneous alternation behavior in the Y maze (*64*). The Y mazes consisted of three arms (4 × 30 × 15 cm) that were the same, with a 120º angle between each of the two arms. Each mouse was placed in one of the Y maze arms and allowed to explore freely through the maze for 5 min. The sequence and total number of arms entered were recorded. An arm entry was considered complete when both hind paws were in the arm. The apparatus was cleaned with water and ethanol between each passage. The percentage of spontaneous alternation was determined by the number of trials containing entries into all three arms/maximum possible alternations (total number of arms entered - 2) × 100.

*Novel object recognition test.* Short-term memory was evaluated by performing a novel object recognition test (*65*). The apparatus consisted of a 60 × 60 × 30 cm acrylic box with white walls and a floor. Animals received 5 min sessions in the empty box for habituation to the apparatus and test room. Then, 24 h later, each mouse was exposed to two familiar objects (round block, 4 cm in diameter) during a 5 min training stage in the box. Next, the animals were placed back in the box and exposed to novel object (rectangle block, 4 × 4 × 4 cm) and the familiar object for another 5 min (test stage), 24 h after the training stage. The time spent exploring each object was measured. The recognition index reflecting the short-term memory ability was calculated as the ratio of time spent exploring the novel object over the total exploration time.

### Immunohistochemistry

For all histological analyses, mice were deeply anesthetized and transcardially perfused with phosphate-buffered saline (PBS) followed by 4% paraformaldehyde (PFA) in 0.2 M phosphate buffer. The brains were removed, post-fixed in 4% PFA, and cryoprotected in 30% sucrose at 4ºC for three days. Brains were divided into 40-μm thick coronal sections on a cryostat microtome (Leica Biosystems, Germany). The brain sections were first incubated for 1.5 h in a blocking solution (0.3% Triton-X, 2% goat serum, and 2% donkey serum in 0.1 M PBS) and then immunostained with a mixture of primary antibodies in a blocking solution at 4°C. Primary antibodies used are as follow: rabbit anti-TH (1:1,000; sc-14007, Santa Cruz, USA), rabbit anti-c-Fos (1:500; sc-253, Santa Cruz), goat anti-pMCH (1:500; sc-14507, Santa Cruz), mouse anti-NeuN (1:1000; MAB377, Millipore), and goat anti-MCH1R (1:500; sc-5534, Santa Cruz). After washing three times in PBS, the sections were incubated in the corresponding fluorescent secondary antibodies for 1 h at room temperature. Then, they were washed with PBS three times. If needed, DAPI (1:3,000, Pierce) staining was performed. Finally, sections were mounted with a fluorescent mounting medium (S3023, Agilent, USA) and dried. A series of fluorescent images were obtained with an A1 Nikon or FV-1000 Olympus confocal microscopes, and Z-stack images in 3-μm steps were processed for further analysis using NIS-Elements (Nikon, Japan) software and ImageJ program (NIH, MD, USA). Any alterations in brightness or contrast were equally applied to the entire image set. Specificity of primary antibody and immunoreaction was confirmed by omitting primary antibodies or changing fluorescent probes of the secondary antibodies.

### DAB staining and stereological quantification

The 30-μm thick coronal sections for SNpc and striatum were immunostained with a DAB peroxidase substrate kit (SK-4100, Vector Laboratories Inc, USA). The sections were blocked in a solution comprised of 0.3% BSA and 3% Triton X-100 for 1 h. Then samples were activated by using anti-TH primary antibody (1:3000, sc-14007, Santa Cruz). Thereafter, sections were incubated with biotinylated secondary antibody (BA-1000, Vector Laboratories Inc., USA) for 1 h. After incubation, sections were activated by avidin-biotinylated peroxidase complex (Vectastain Elite ABC kit, Vector Laboratories Inc.) solution for 1.5 h. Sections were then developed with DAB peroxidase substrate kit. Finally, sections were mounted with mounting solution and dried. A series of bright field images of $TH^+$ cells and fibers in the SNpc and striatum were obtained by using a microscope (BX53, Olympus, Japan).

An unbiased three-dimensional counting method, stereological estimation of the total number of $TH^+$ neurons in the SNpc area was performed using the optical fractionator method using Stereo Investigator 11 (11.01.2 64-bit, MBF Bioscience, USA). For stereological purposes, unilateral sets of 1/6 section, ~8 systematically random sections, of about 240 μm apart, were taken from the brain spanning in all mice. The counted sections covered the rostral tip of the SNpc. An unbiased counting frame of known area (48 × 36 μm = 1,728 $\mu m^2$) was placed randomly on the first counting area and systematically moved through all counting areas (166.2 × 111.12 μm = 18,468 $\mu m^2$) until the entire delineated area was sampled. Counting was performed using a low magnification objective lens (×10). The estimated total number of positive neurons was calculated according to the optical fractionator formula.

### In-vivo microendoscopy

*Imaging.* A microendoscope (nVista 2.0, Inscopix, USA) was used to record calcium signals from MCH neurons within LH in mice. To observe calcium activities of MCH cells, *Ai148* transgenic mice were injected with 1.5 μL of *AAV-pMCH-Cre* virus diluted by 1:5 (resulting titer: 1.14 x10^13 GC/mL) into the left LH. Two weeks after virus injection, mice were implanted with GRIN lenses (ProView™ Integrated Lens - 0.5 mm diameter, 8.4 mm length, Inscopix, USA) with attached magnetic bases able to fix the microendoscope. Lens tips were fixed in LH (AP: -1.0 mm, ML: -1.2 mm from bregma, DV: -4.8 mm from skull surface). After confirming that mice showed calcium activity, mice were anesthetized by 3% isoflurane, and then maintained at 1% isoflurane for the light anesthesia during the experiment session. A typical session involved approximately 10 minutes of calcium signal acquisition using Inscopix Data Acquisition Software at 20 frames per second and simultaneous acupuncture stimuli, both with and without lidocaine administration. Excitation laser power and digital gain were customized for each mouse according to the conditions of its cell population.

*Processing.* Raw data obtained were processed with Mosaic data processing software (Inscopix Data Processing 1.2.0, Inscopix, USA), and, first underwent motion correction to control potential motion artefacts. Next, the software normalized the brightness every pixel value in the raw video, thus generating each pixel's deviation trace, $dF/F_0$, from the mean baseline. An inbuilt Principle Component Analysis and Independent Component Analysis algorithm automatically generated cell candidates based on pixels exhibiting synchronized spatial and temporal brightness changes. The resulting regions of interest representing candidate neurons were further filtered manually based on cell shape and signal-to-noise ratio. Resulting cell calcium activity traces were aligned with the experiment footage.

### SHIELD tissue processing and light-sheet microscopy

Mice were transcardially perfused with ice-cold PBS and then with the SHIELD perfusion solution (PCK-500, Passive clearing kit, Lifecanvas, USA) as described previously (*66*, *67*). Dissected brains were incubated in the same perfusion solution at 4°C for 48 h. Tissues were then transferred to the SHIELD-OFF solution and incubated at 4°C for 24 h. Following the SHIELD-OFF step, brains were placed in the SHIELD-ON solution and incubated at 37 °C for 24 h. SHIELD-fixed brains were cleared passively for a couple of weeks at 45 °C in a buffer solution. For optical clearing, delipidated tissues were incubated in a solution of 50% EasyIndex diluted in PBS overnight at 37°C. The solution was then changed to 100% EasyIndex and incubated overnight at

37°C. Tissues became transparent without any visible haze at the tissue–medium interface. 3D light-sheet images were taken by Zeiss Light sheet Fluorescence microscopy (LSFM) 7.

Slice electrophysiology

*Acute brain slice preparation.* Mice were anesthetized using vaporized isoflurane. Brain was quickly removed and immersed in an ice-cold cutting solution that contained (in mM): 130 NaCl, 24 $NaHCO_3$, 3.5 KCl, 1.25 $NaH_2PO_4$, 3.0 $MgCl_2$, 1.0 $CaCl_2$ and 10 d-glucose, pH 7.4. All the solution was gassed with 95% $O_2$ and 5% $CO_2$. After trimming the hemisected cerebrum, 300-μm-thick horizontal slices were cut using vibrating microtome (Neo LinearSlicer MT, DSK, Japan) and transferred to an artificial cerebrospinal fluid (aCSF) recording solution (in mM): 130 NaCl, 24 $NaHCO_3$, 3.5 KCl, 1.25 $NaH_2PO_4$, 1.5 $MgCl_2$, 1.5 $CaCl_2$ and 10 d-(+)- glucose, pH 7.4. Slices were incubated at room temperature for 1 h before recording.

*Action potential recording.* Slices were transferred to a recording chamber that was perfused with aSCF solution. The slice chamber was mounted on the stage of an upright Olympus microscope and viewed with a 60x objective lens. Cellular morphology was visualized by charge-coupled device camera and the Imaging Workbench software (INDEC BioSystems). $GFP^+$ or $mCherry^+$ $MCH^{LH/ZI}$ neurons ($MCH^{LH\rightarrow HPC}$ and $MCH^{LH/ZI\rightarrow SNpc}$ neurons, respectively) were patched with whole-cell configuration. The holding potential was −60 mV. Pipette resistance was 6-8 MΩ and filled with an internal solution (in mM): 140 K-gluconate, 10 hydroxyethyl piperazine ethane sulfonic acid, 7 NaCl, 0.5 ethylene glycol tetraacetic acid and 2 Mg-ATP adjusted to pH 7.4 in current-clamp mode. Current step was given from -120 pA to 80 pA with 20-pA intervals. Electrical signals were digitized and sampled at 10 kHz with Digidata 1322A (Axon Instruments) and Multiclamp 700B amplifier (Molecular Devices) using pCLAMP 10.2 software (Molecular Devices). Raw data were low-pass filtered at 2 kHz and collected for off-line analysis at a sampling rate of 10 kHz using pClamp 10.2 software.

*Long-term potentiation.* Acute brain slices were prepared as aforementioned with slight modification. The 350-μm-thick horizontal hippocampal slices were prepared and incubated at room temperature for 1 h before recording. fEPSP was recorded from CA1 stratum radiatum of the hippocampus using a glass pipette filled with aCSF (1-3 MΩ) upon electrical stimulation triggered by a concentric bipolar electrode (CBBPE75, FHC, USA) placed in the Schaffer collateral pathway. Evoked fEPSP responses were digitized and sampled at 10 kHz with Digidata 1322A (Axon Instruments) and Multiclamp 700B amplifier (Molecular Devices) using pCLAMP 10.2 software (Molecular Devices). The stimulation intensity was adjusted to obtain fEPSP slopes of 40-50 % to the maximum. Basal fEPSP response was monitored by electrical stimulations at 0.1 Hz. Theta-burst stimulation (TBS) for inducing LTP consisted of three trains (1.2 s) of five pulses at 100 Hz and pulse with was 0.2 ms with an inter-burst interval of 200 ms.

Ex-vivo calcium imaging

300-μm-thick hippocampal slices were acutely prepared as aforementioned. Imaging was acquired at one frame per second with a 60x water-immersion objective lens, and a 488-nm fluorescent imaging filter was utilized for GCaMP6f imaging. CNO (5 μM) was applied for 100 sec after baseline stabilization. Fluorescence imaging was acquired and analyzed with Imaging Workbench (Indec Biosystems), and analyzed with ImageJ software (NIH).

RNA sequencing

*RNA preparation.* For RNA sequencing, we harvested SNpc and hippocampal tissues from each group. We extracted total RNA using the easy-spin™ Total RNA Extraction Kit (#17221, iNtRON, Korea) and assessed the RNA concentration with Nanodrop one (Thermo scientific). Sample libraries for sequencing were prepared by the Novaseq Reagent Kit Preparation Guide (Illumina, USA) by Macrogen Inc. (Korea).

*mRNA-Seq Data.* We pre-processed the raw reads from the sequencer to remove low quality and adapter sequence before analysis and aligned the processed reads to the *Mus musculus (mm10)* using HISAT v2.1.0(1). HISAT utilizes two types of indexes for alignment (a global, whole-genome index and tens of thousands of small local indexes). These two types of indexes are constructed using the same BWT (Burrows–Wheeler transform) a graph FM index (GFM) as Bowtie2. Because of its use of these efficient data structures and algorithms, HISAT generates spliced alignments several times faster than Bowtie and BWA widely used. The reference genome sequence of *Mus musculus (mm10)* and annotation data were downloaded from the NCBI. Transcript assembly and abundance estimation using StringTie(2, 3). After alignment, StringTie v2.1.3b was used to assemble aligned reads into transcripts and to estimate their abundance. It provides the relative abundance estimates as Read Count values of transcript and gene expressed in each sample.

*Statistical analysis of gene expression level.* The relative abundances of gene were measured in Read Count using StringTie. We performed the statistical analysis to find differentially expressed genes using the estimates of abundances for each gene in samples. Genes with one more than zeroed Read Count values in the samples were excluded. To facilitate log2 transformation, 1 was added to each Read Count value of filtered genes. Filtered data were log2-transformed and subjected to RLE normalization. Statistical significance of the differential expression data was determined using nbinomWaldTest using DESeq2 and fold change in which the null hypothesis was that no difference exists among groups. False discovery rate (FDR) was controlled by adjusting p value using Benjamini-Hochberg algorithm. For DEG set, hierarchical clustering analysis was performed using complete linkage and Euclidean distance as a measure of similarity. The criteria for DEG was FPKM>=1(in more than one group), $|\text{fold change}| \geq 1.5$, and adjusted $p < 0.05$ (for SNpc) or raw $p < 0.05$ (for HPC). GO analysis was performed based on Gene Set Enrichment Analysis (GSEA; http://www.gsea-msigdb.org/gsea/msigdb/mouse/annotate.jsp) with FDR q value < 0.05.

MCHR1 shRNA synthesis

For gene-silencing Mchr1, we chose three candidate target sequences from coding sequence of mouse Mchr1 (NM_145132) gene: **1)** 5’-GCA CAA GGA GTG TCT CCT ACA-3’, **2)** 5’-GCA ACG TCC CTG ACA TCT TCA-3’, **3)** 5’-GCC TCA ATC CCT TTG TGT ACA-3’. We prepared MCHR1-shRNA candidates whose sequences of complementary oligomers were as follows: **1)** 5’-TGC ACA AGG AGT GTC TCC TAC ATT CAA GAG ATG TAG GAG ACA CTC CTT GTG CTT TTT TC-3’ and 3’-TCG AGA AAA AAG CAC AAG GAG TGT CTC CTA CAT CTC TTG AAT GTA GGA GAC ACT CCT TGT GCA-5’, **2)** 5’-TGC AAC GTC CCT GAC ATC TTC ATT CAA GAG ATG AAG ATG TCA GGG ACG TTG CTT TTT TC-3’ and 3’-TCG AGA AAA AAG CAA CGT CCC TGA CAT CTT CAT CTC TTG AAT GAA GAT GTC AGG GAC GTT

GCA-5’, and **3)** 5’-TGC CTC AAT CCC TTT GTG TAC ATT CAA GAG ATG TAC ACA AAG GGA TTG AGG CTT TTT TC-3’ and 3’-TCG AGA AAA AAG CCT CAA TCC CTT TGT GTA CAT CTC TTG AAT GTA CAC AAA GGG ATT GAG GCA-5’ (Table S3). To test the knockdown efficiency of MCHR1-shRNA candidates, we obtained GFP-tagged Mchr1 (NM_145132) ORF clone from Origene (#MG219644). The knockdown efficiency was tested by quantitative RT-PCR (qPCR) with cDNA from HEK293T cell line (Korean Cell Line Bank) which were transfected with the GFP-tagged Mchr1 full clone and shRNA vectors. qPCR was performed with QuantStudio 1 Real-Time PCR machine (Applied Biosystems, USA). qPCR was carried out using SYBR Green PCR Master Mix (Applied Biosystems). In brief, reactions were performed in duplicates in a total volume of 10 ml containing 10 pM primer, 40 ng cDNA, and 5 ml power SYBR Green PCR Master Mix. The mRNA level was normalized to that of GAPDH mRNA. Fold-induction was calculated using the $2^{-\Delta\Delta Ct}$ method.

Preparation of human dopaminergic (hDA) neurons from human iPSCs

The hDA neurons were derived from human iPSCs (Coriell, GM25256) by using STEMdiff™ Midbrain Neuron Differentiation Kit (STEM CELL tech, #100-0038) according to manufacturer instructions. Immunocytochemistry was performed for counting the TH- and MAP2-positive cell numbers. qPCR was carried out for measuring the mRNA expression levels of TH, MAP2, TUJ-1, and GAP43 (Table S3).

Quantification and statistical analysis

Statistical analyses were performed using Prism 9 (GraphPad Software, Inc.). Differences between two different groups were analyzed with the two-tailed Student’s unpaired t-test. For comparison of multiple groups, one-way analysis of variance (ANOVA) with Tukey’s or Dunnett’s multiple comparison test, or two-way ANOVA with Bonferroni’s multiple comparison test was assessed. For assessment of change of a group by a certain intervention, the significance of data was assessed by the two-tailed Student’s paired t-test or repeated measure one-way ANOVA. The normality of the distribution of each dataset was tested. When the data was not normally distributed, we performed appropriate non-parametric tests, such as Mann-Whitney test or Kruskal-Wallis ANOVA test. For comparisons of two or multiple groups, we also tested if the variances were statistically different across the groups. If the variance was different, appropriate corrections were applied to the statistical tests. $P < 0.05$ was considered to indicate statistical significance throughout the study. The significance level is represented as asterisks ($^{*}P < 0.05$, $^{**}P < 0.01$, $^{***}P < 0.001$; *NS*, not significant). Unless otherwise specified, all data are presented as mean ± SEM. No statistical method was used to predetermine sample size. Sample sizes were determined empirically based on our previous experiences or the review of similar experiments in literature. The numbers of animals used are described in the corresponding figure legends or on each graph. All experiments were done with at least three biological replicates. Experimental groups were balanced in terms of animal age, sex, and weight. Animals were genotyped before experiments, and they were all caged together and treated in the same way. Prior to acupuncture stimulation, virus injection, or drug administration, animals were randomly and evenly allocated to each experimental group. The data analysis of animal experiments was performed by two independent investigators. However, investigators were not blinded to outcome assessments.

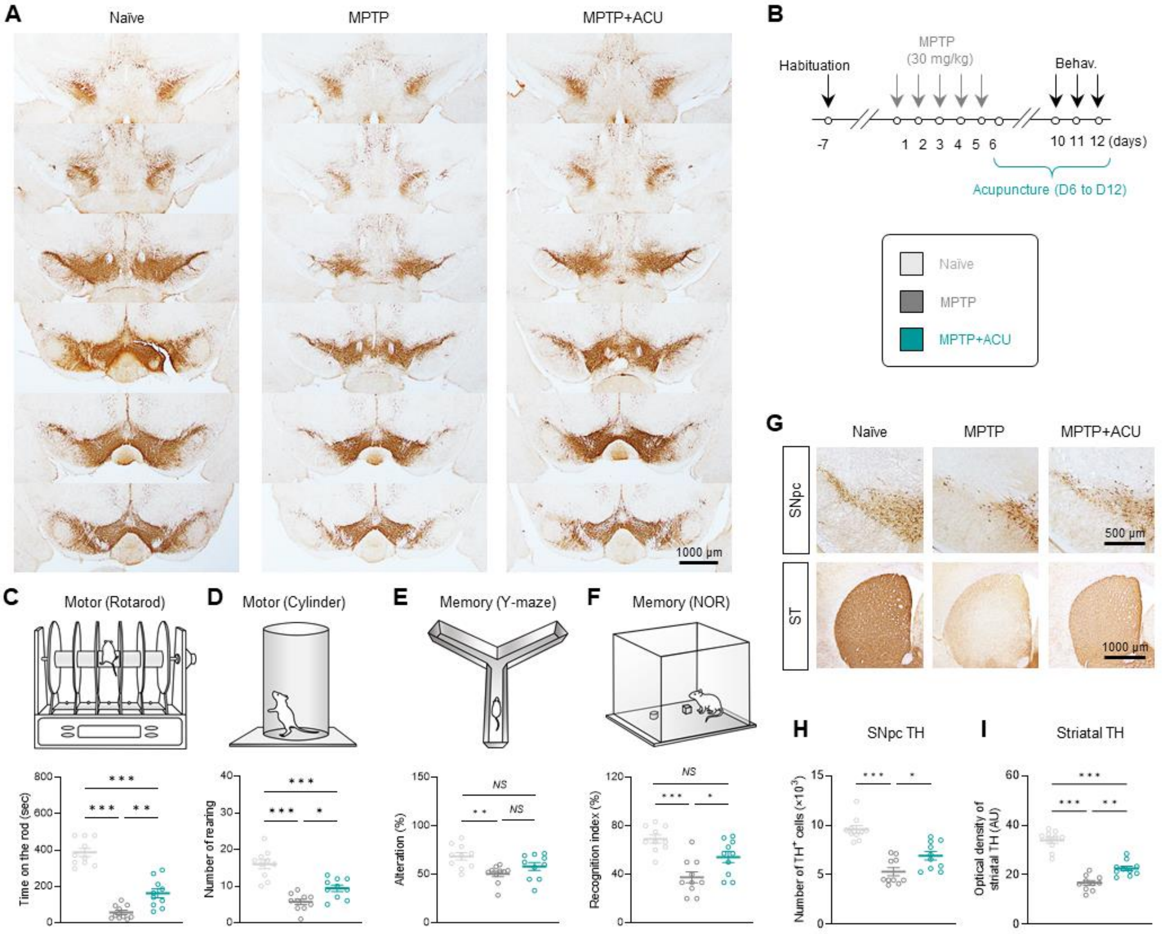


**Fig. S1. Representative images of serially sectioned SNpc tissues stained with TH antibody and therapeutic effects of acupuncture starting after MPTP treatment.** (A) TH-positive neurons from the rostral to the caudal portions of the SNpc. (B) Timeline of experiments to investigate the therapeutic effect of acupuncture which started after MPTP administration (post-ACU). (C-F) Assessment of motor and memory functions. Post-ACU treatment for one week after induction of the MPTP model has a therapeutic effect for motor dysfunction (One-way ANOVA, n = 10 per group; for rotarod: $F_{2, 27} = 66.2$, $P < 0.001$; for cylinder: $F_{2, 27} = 29.6$, $P < 0.001$; *post-hoc* Tukey's test: $^{***}P < 0.001$, $^{**}P < 0.01$, $^{*}P < 0.05$) and memory impairment (Kruskal-Wallis ANOVA for Y-maze, n = 10 per group: $H_3 = 9.547$, $P = 0.008$; One-way ANOVA for NOR, n = 10 per group: $F_{2, 27} = 14.1$, $P < 0.001$; *post-hoc* Tukey's test: $^{***}P < 0.001$, $^{**}P < 0.01$, $^{*}P < 0.05$, *NS*, not significant). (G) Representative images of TH expression in SNpc and striatum in MPTP model with or without post-ACU. (H - I) Quantification of the number of TH-positive neurons and optical density of striatal TH expression. Post-ACU treatment partially but significantly restored the expression of TH level in the SNpc and striatum (H, Kruskal-Wallis ANOVA for SNpc, n = 10 per group: $H_3 = 20.20$, $P < 0.001$; I, One-way ANOVA for striatum, n = 10 per group: $F_{2, 27} = 69.7$, $P < 0.001$; *post-hoc* Tukey's test: $^{***}P < 0.001$, $^{**}P < 0.01$, $^{*}P < 0.05$). Data are shown as mean ± SEM.

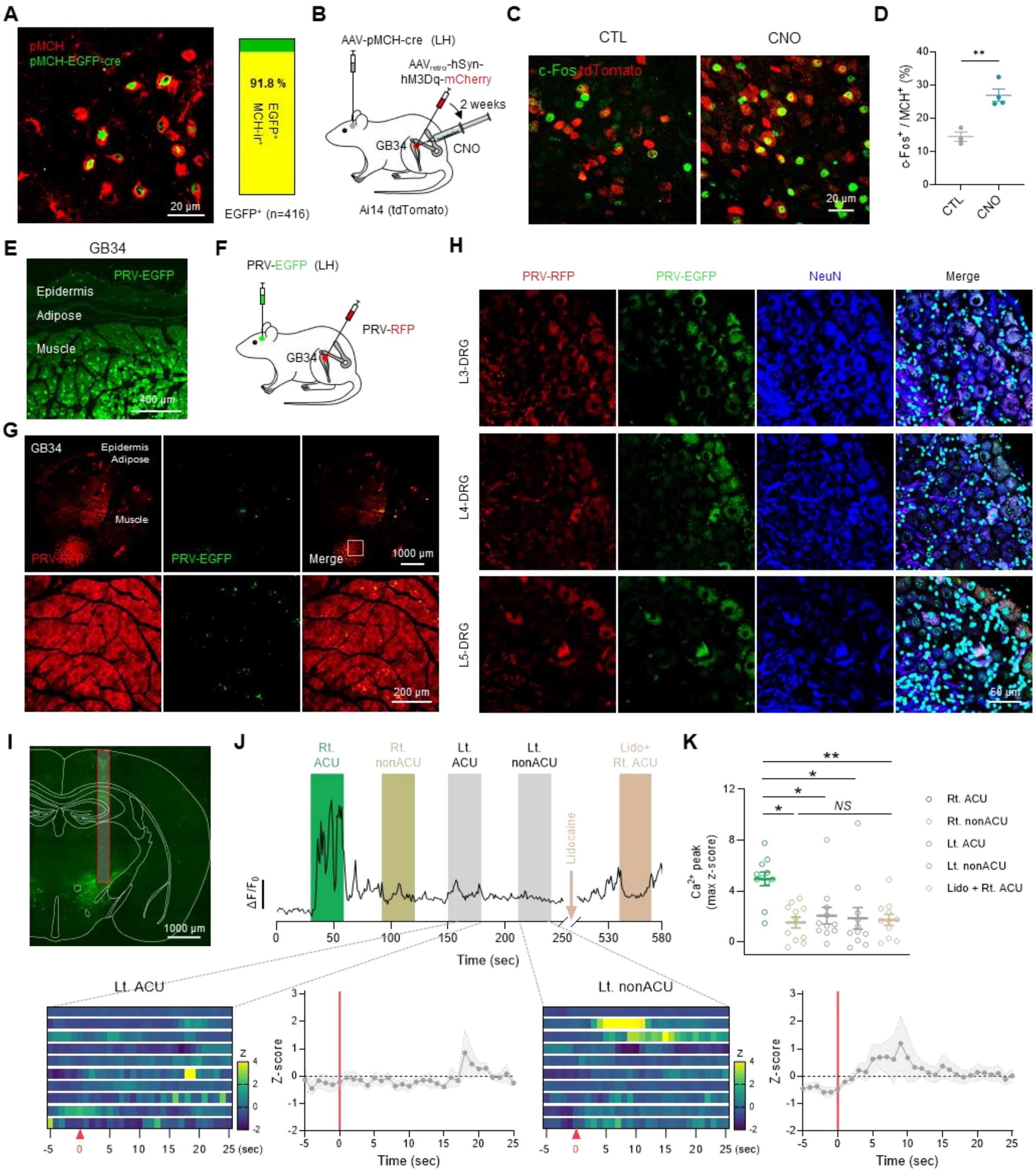


**Fig. S2. The trans-synaptic retrograde labeling for visualization of neural pathways from hindlimb acupoint GB34 to the LH/ZI.** (A) Left, the representative confocal image of pMCH-immunoreactivity and EGFP which is expressed by $AAV_{DJ}$-pMCH-EGFP-cre virus injection into the mouse LH. Right, Quantification of co-expression of pMCH-EGFP-cre and pMCH (N = 4 mice, n = 416 cells) reveals that EGFP expression was specifically restricted to pMCH-immunolabeled neurons (91.8%, n = 335 cells). (B) Schematic diagram of chemogenetic stimulation of peripheral afferent nerve fibers at acupoint GB34. (C) Representative confocal images of c-Fos expression in the $MCH^{LH/ZI}$ neurons 1 h after chemogenetic stimulation of acupoint GB34. (D) Quantification of c-$Fos^+$ $MCH^{LH/ZI}$ neurons upon chemogenetic stimulation of acupoint GB34 (two-tailed unpaired t-test, n = 3-4 per group; $t_5 = 4.936$, $P = 0.0043$). (E) Representative confocal image of the GB34 injected with PRV-CMV-EGFP. (F) Schematic diagram depicting the injection strategy with double retrograde PRVs. PRV-CMV-RFP and PRV-CMV-EGFP were injected into the GB34 and LH, respectively. (G) Representative confocal images of the cross section of skin and muscle at GB34. Because PRV-CMV-RFP is injected into this point, RFP signal is abundantly observed. Notable, EGFP-positive nerve endings which should originated from LH where PRV-CMV-EGFP is injected. (H) Representative confocal images of DRG (L3 to L5) demonstrating double labeled neurons with PRV-RFP and PRV-EGFP. (I) GRIN lens position and GCaMP6f expression in the LH. (J) Top, a representative trace of $Ca^{2+}$ signal of a pMCH neuron upon various acupuncture stimulations. Bottom, heatmaps displaying $Ca^{2+}$ signal of each pMCH neuron and averaged traces of $Ca^{2+}$ signal of pMCH neurons. (K) Quantification of $Ca^{2+}$ peak displayed by a scatter plot. The peak $Ca^{2+}$ signals were significantly increased only by the acupuncture treatment at right GB34. (Kruskal-Wallis ANOVA, n = 11 per group; $H_5 = 16.23$, $P = 0.003$; *post-hoc* Dunnett's test: $^{**}P < 0.01$, $^{*}P < 0.05$, *NS*, not significant). Data are shown as mean ± SEM.

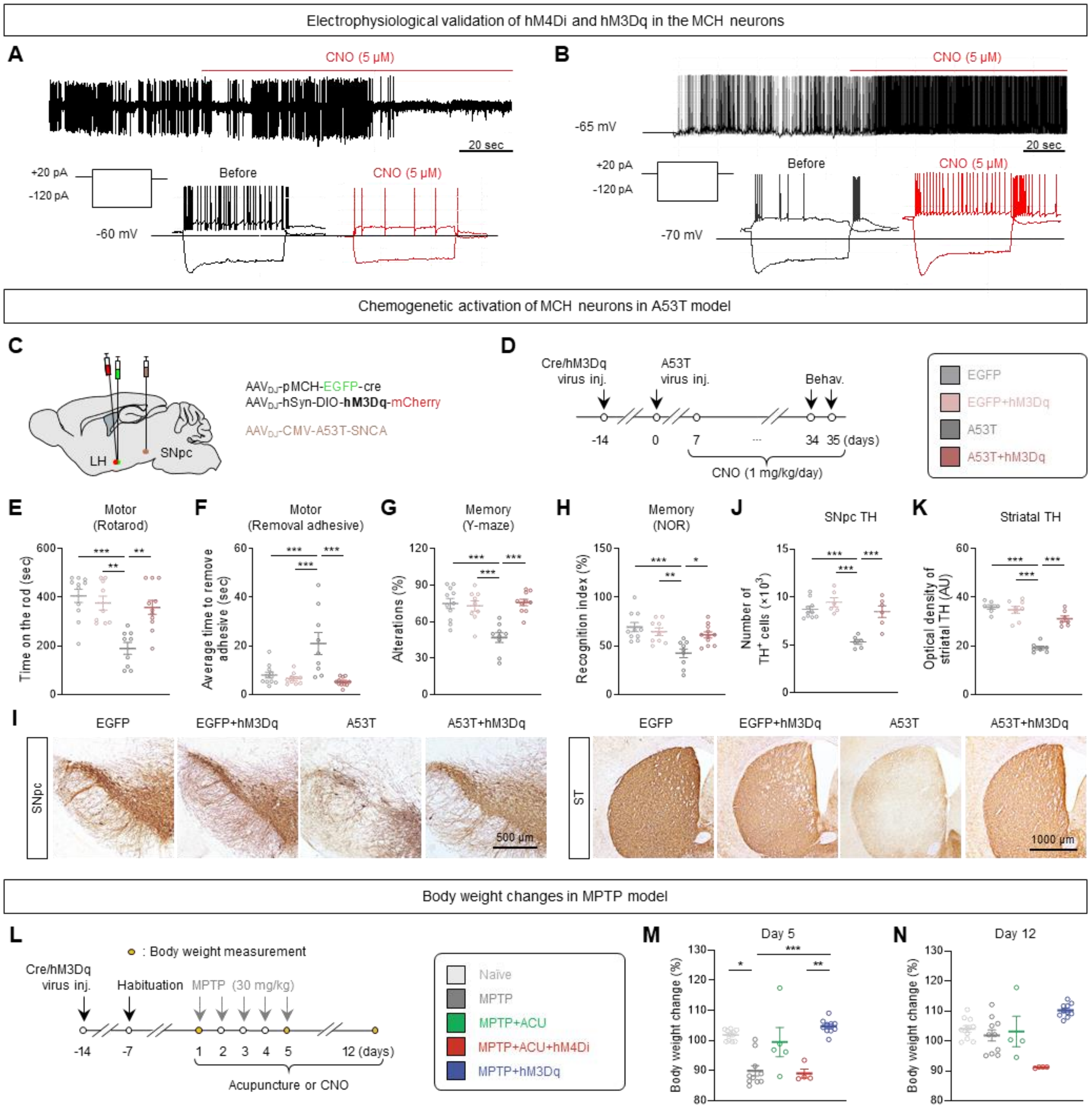


**Fig. S3. Activation of MCH neurons alleviates parkinsonian motor and memory deficits in two different PD mouse models.** (A) Electrophysiological validation of hM3Dq. Top, cell-attached patch clamp recording of spontaneous action potential before and after bath application of CNO (5 μM). Bottom, current clamp recording of action potential induced by current injection (+20 pA). (B) Electrophysiological validation of hM4Di. Top, current clamp recording of spontaneous firing. Bottom, current clamp recording of action potential induced by current injection (+20 pA). (C) Schematic diagram depicting viral strategy for testing chemogenetic approach in A53T model. (D) Experimental timeline of chemogenetic activation of MCH neurons in A53T model. (E-H) Assessment of motor and memory function by rotarod test, adhesive removal test, Y-maze test, and novel object recognition test. Chemogenetic activation of $pMCH^{LH/ZI}$ neurons alleviated the motor dysfunction (E, F; Kruskal-Wallis ANOVA for rotarod: $H_4 = 17.92$, $P < 0.001$; One-way ANOVA for removal adhesive: $F_{3, 36} = 10.4$, $P < 0.001$; *post-hoc* Tukey's test: $^{***}P < 0.001$, $^{**}P < 0.01$, $^{*}P < 0.05$) and memory deficits (G, H; One-way

ANOVA; for Y-maze: $F_{3, 35} = 12.7$, $P < 0.001$; for NOR: $F_{3, 35} = 8.10$, $P < 0.001$; *post-hoc* Tukey's test: $^{***}P < 0.001$, $^{**}P < 0.01$, $^{*}P < 0.05$) in A53T model. (I) Representative images of TH staining in the SNpc and striatum. (J-K)  Quantification of the number of TH-positive neurons and optical density of striatal TH expression. The number of TH-positive cells in the SNpc and the optical density of TH-positive dopaminergic fibers in the striatum were restored by chemogenetic activation of $pMCH^{LH/ZI}$ neurons in the A53T model (One-way ANOVA; for SNpc: $F_{3, 24} = 17.5$, $P < 0.001$; for striatum: $F_{3, 25} = 46.4$, $P < 0.001$; *post-hoc* Tukey's test: $^{***}P < 0.001$). l, Timeline of experiments for MPTP model. m,n, Acupuncture and chemogenetic activation of MCH neurons reverses MPTP-induced body weight loss at day 5 (Kruskal-Wallis ANOVA; for day 5: $H_5 = 26.60$, $P < 0.001$; for day 12: $H_5 = 20.42$, $P < 0.001$; *post-hoc* Tukey's test: $^{***}P < 0.001$, $^{**}P < 0.01$, $^{*}P < 0.05$). Data are shown as mean ± SEM.

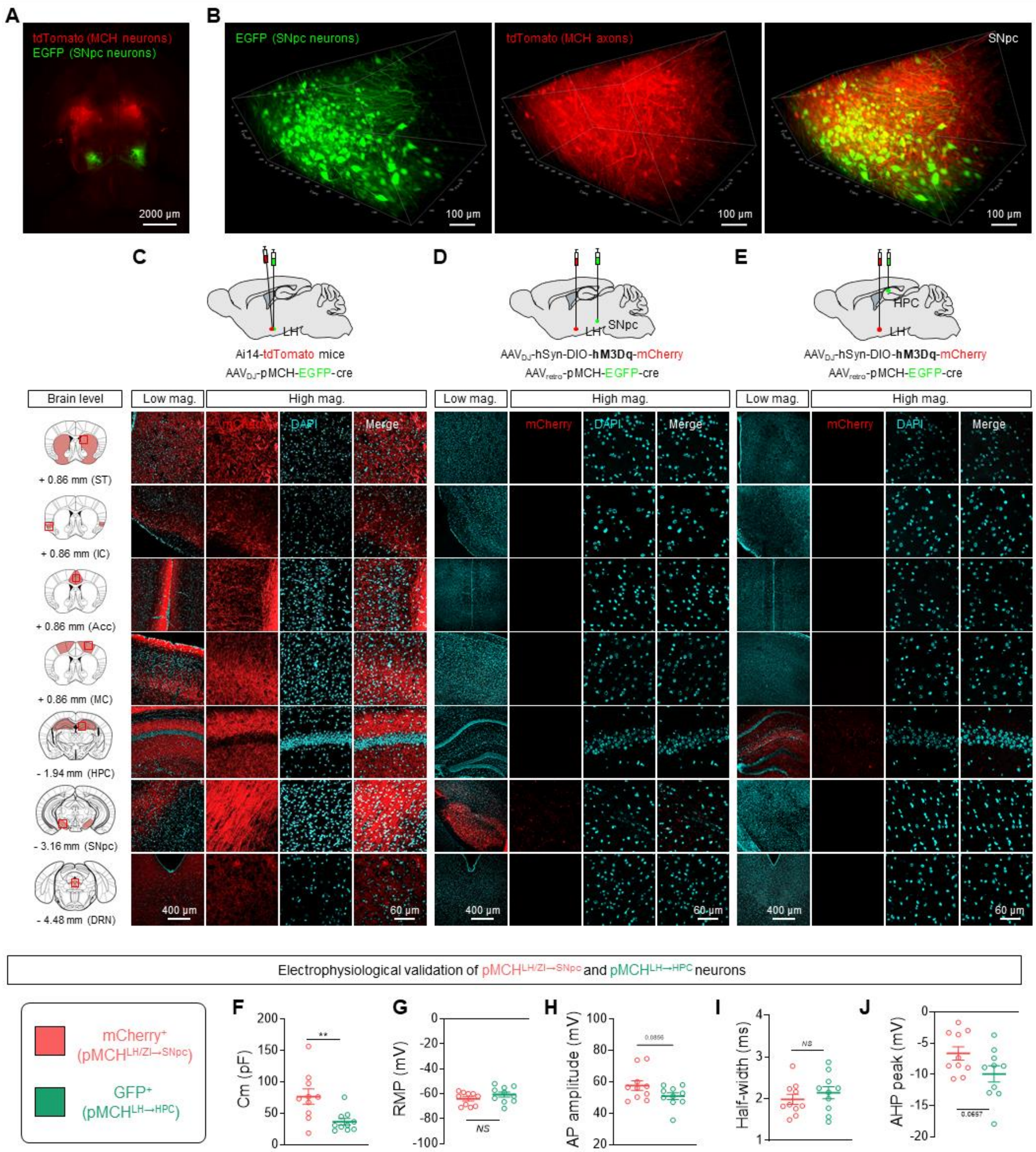


**Fig. S4. Anatomical analysis of MCH neuronal projections and Intrinsic electrophysiological properties of MCH$^{LH/ZI\rightarrow SNpc}$ and MCH$^{LH\rightarrow HPC}$ neurons.** (A) Three-dimensional rendering of a cleared mouse brain showing brain-wide injection patterns of MCH neurons labeled by tdTomato and EGFP (SNpc neurons) in the Ai14 (Rosa26-Stop-tdTomato) mouse. (B) SNpc neurons (EGFP) co-expressed with axons of MCH neurons projected from LH (tdTomato) in the SNpc. (C-E) Confocal images of striatum (ST), insular cortex (IC), anterior cingulate cortex (ACC), motor cortex (MC), hippocampus (HPC), substantia nigra pars compacta

(SNpc), dorsal raphe nucleus (DRN) in the mice of tdTomato-labeling within $MCH^{LH/ZI}$ neurons (C), $MCH^{LH/ZI \rightarrow SNpc}$ neurons (D), and $MCH^{LH \rightarrow HPC}$ neurons (E). (F) Membrane capacitance (two-tailed unpaired t-test, n = 10 per group; $t_{18} = 3.053$, $P = 0.007$). (G) Resting membrane potential (two-tailed unpaired t-test, n = 10 per group; $t_{18} = 1.303$, $P = 0.2088$). (H) AP amplitude (two-tailed unpaired t-test, n = 10 per group; $t_{18} = 1.819$, $P = 0.0856$). (I) AP threshold AP half-width (two-tailed unpaired t-test, n = 10 per group; $t_{18} = 0.8436$, $P = 0.4100$). (J) After-hyperpolarization peak (two-tailed unpaired t-test, n = 10 per group; $t_{18} = 1.960$, $P = 0.0657$)

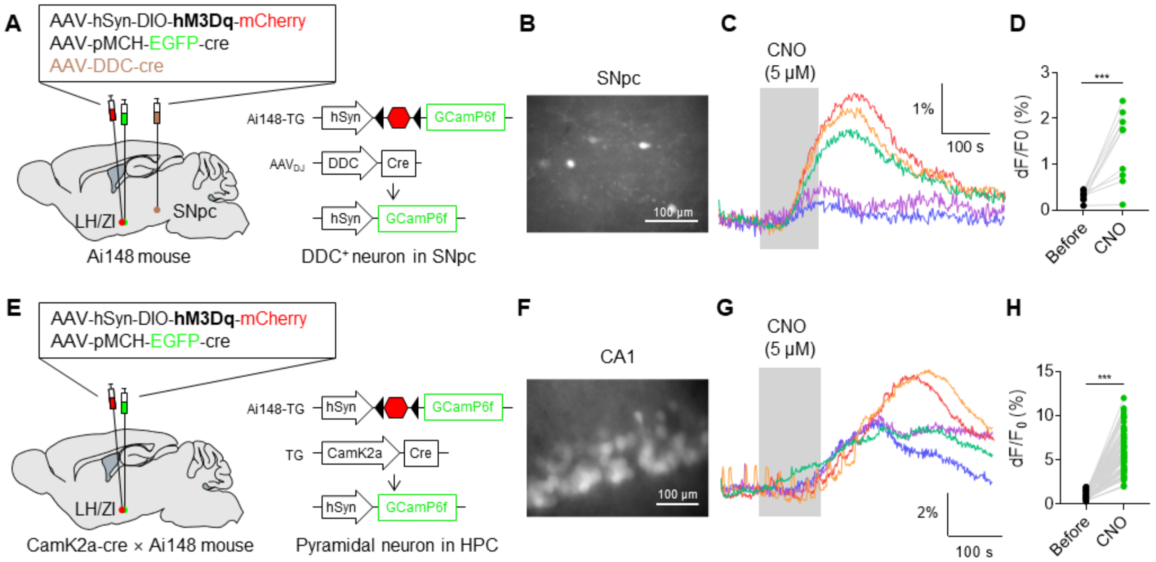


**Fig. S5. Functional connection of the discrete neural projections of MCH neurons from LH and ZI to SNpc and HPC.** (A and E) Schematic diagram of *ex vivo* $Ca^{2+}$ imaging of SNpc dopaminergic neurons (A) and CA1 hippocampal pyramidal neurons (E) upon hM3Dq-mediated chemogenetic activation of $MCH^{LH/ZI}$ neurons. (B and F) Representative images of GCaMP6f expression in SNpc (B) and CA1 HPC (F). (C and G) $Ca^{2+}$ signal traces of SNpc dopaminergic neurons (C) and CA1 hippocampal pyramidal neurons (G). (D and H) Quantification of $Ca^{2+}$ peak upon chemogenetic activation of MCH neurons in each region (two-tailed unpaired t-test; SNpc, n = 10 per group, $t_{18} = 4.593$, $P < 0.001$; CA1, n = 101 per group, $t_{200} = 21.52$, $P < 0.001$).

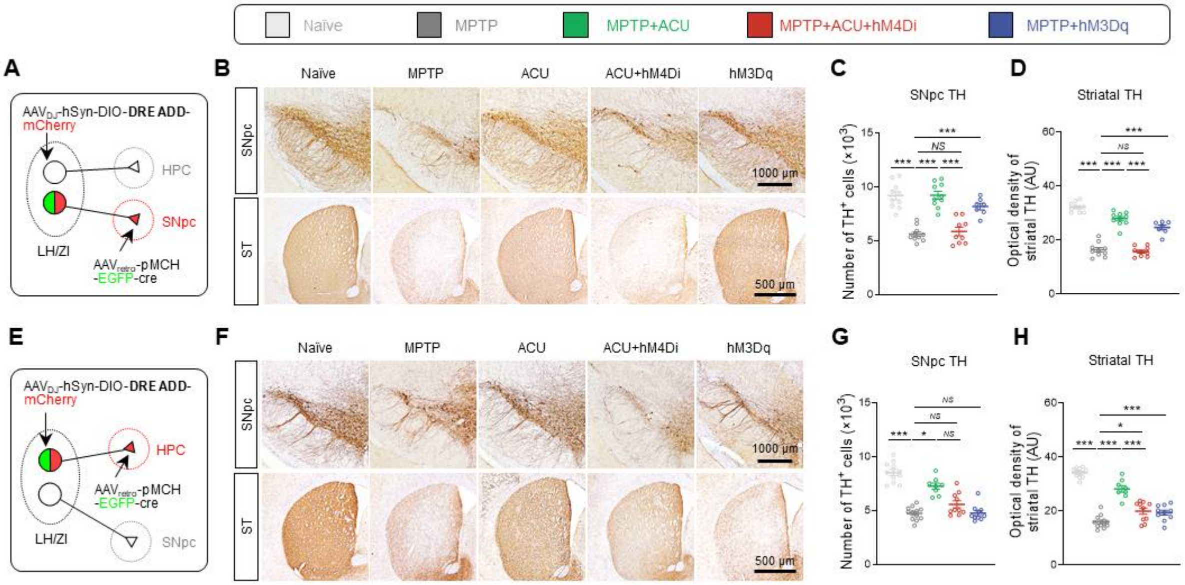


**Fig. S6. MCH$^{LH/ZI\rightarrow SNpc}$ projections, but not MCH$^{LH\rightarrow HPC}$ projections, are involved in the recovery of dopaminergic neurons.** (A) Schematic of viral strategy for projection-specific chemogenetic manipulation of MCH$^{LH/ZI\rightarrow SNpc}$. (B) Representative images of TH expression in the SNpc and striatum. (C) Numbers of TH-positive dopaminergic neurons in the SNpc (One-way ANOVA, n = 7-10 per group: $F_{4,\,41}$ = 28.7, $P$ = 0.465; *post-hoc* Tukey's test: $^{***}P < 0.001$, *NS*, not significant). (D) Quantification of optical density of striatal TH (One-way ANOVA, n = 7-10 per group: $F_{4,\,41}$ = 99.9, $P$ = 0.756; *post-hoc* Tukey's test: $^{***}P < 0.001$, *NS*, not significant). (E) Schematic of viral strategy for projection-specific chemogenetic manipulation of MCH$^{LH\rightarrow HPC}$. (F) Representative images of TH expression in the SNpc and striatum. (G) Numbers of TH-positive dopaminergic neurons in the SNpc (One-way ANOVA, n = 8-12 per group: $F_{4,\,46}$ = 42.3, $P$ = 0.542; *post-hoc* Tukey's test: $^{***}P < 0.001$, $^{*}P < 0.05$, *NS*, not significant). (H) Quantification of optical density of striatal TH (One-way ANOVA, n = 8-12 per group: $F_{4,\,46}$ = 74.4, $P$ = 0.251; *post-hoc* Tukey's test: $^{***}P < 0.001$, $^{*}P < 0.05$). Data are shown as mean ± SEM.

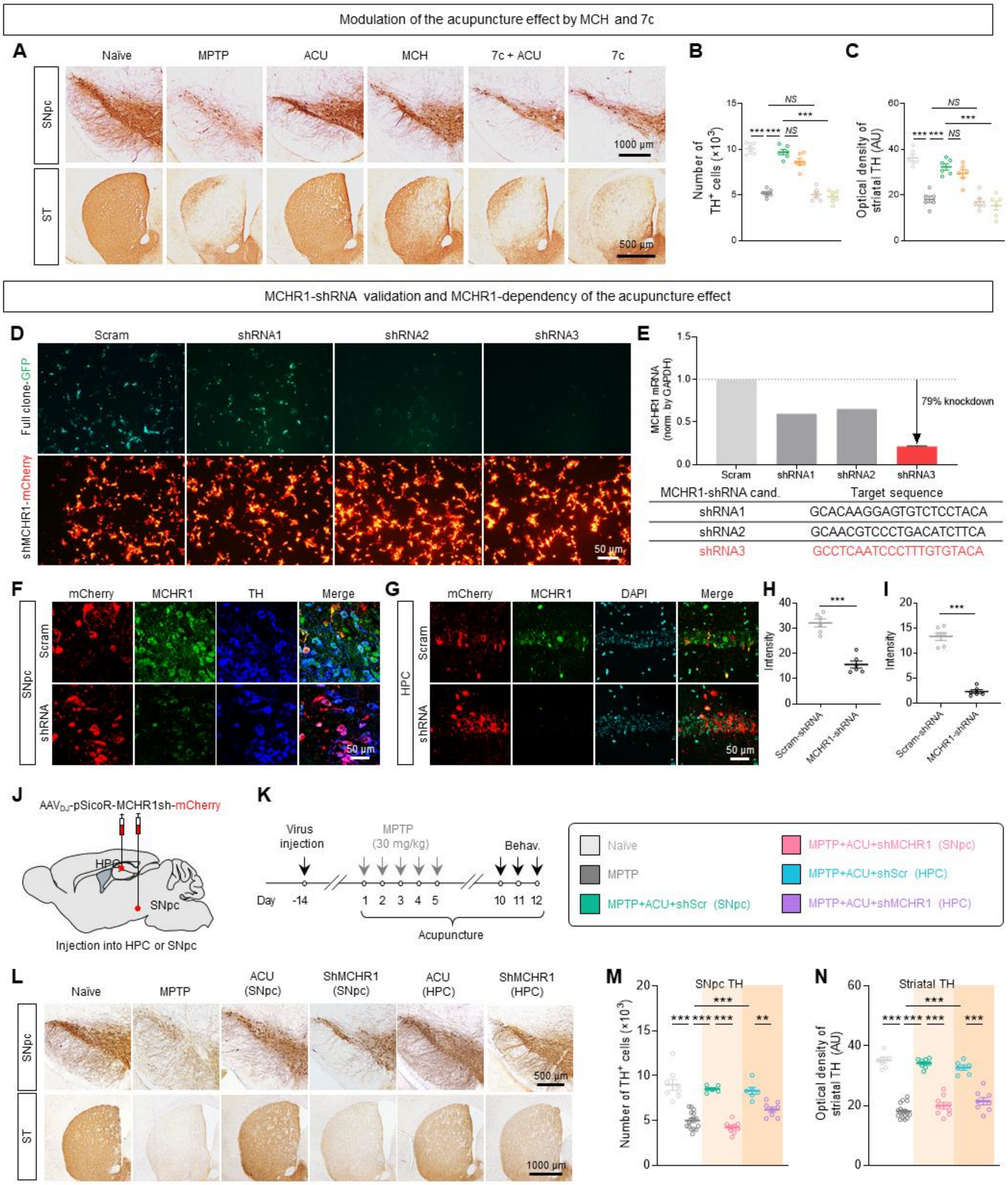


**Fig. S7. MCHR1 gene-silencing blocks the acupuncture effect.** (A) Representative images of TH expression in the SNpc and striatum. (B) Numbers of TH-positive dopaminergic neurons in the SNpc (One-way ANOVA, n = 6 per group: $F_{5,\,30} = 78.9$, $P = 0.855$; *post-hoc* Tukey's test: $^{***}P < 0.001$, *NS*, not significant). (C) Quantification of optical density of striatal TH (One-way ANOVA, n = 6 per group: $F_{5,\,30} = 32.4$, $P = 0.962$; *post-hoc* Tukey's test: $^{***}P < 0.001$, *NS*, not significant). (D) Representative fluorescent images of HEK293T cells displaying the expression of shRNA candidates (mCherry) and the reduced expression of MCHR1 full clone (GFP) 24 h after co-transfection of MCHR1 full clone and the shRNA candidates. (E) Top, *in vitro*

knockdown efficiency of MCHR1-shRNA candidates. Relative levels of MCHR1 mRNA expression were quantified by normalizing with GAPDH mRNA. Knockdown efficacy was most pronounced by transfection with shRNA3, showing ~79% decrease compared to Scram non-knockdown control. Bottom, the target sequences of shRNA candidates. (F-I)**,** In vivo knockdown efficiency of MCHR1-shRNA3 in SNpc and HPC, validated by immunohistochemistry. Compared with the Scrambled shRNA, shRNA significantly reduced the intensity of MCHR1 expression in the SNpc (H) and HPC (I) (two-tailed unpaired t-test, n = 6 per group; for SNpc: $t_{10} = 7.782$; $P < 0.0001$; for HPC: $t_{10} = 13.07$; $P < 0.0001$). (J) Schematic diagram of the location of the $AAV_{DJ}$-pSicoR-MCHR1sh-mCherry virus injection in the SNpc and HPC. (K**)** Timeline of experiments for *in vivo* gene-silencing of MCHR1 in the MPTP model. (L) Representative images of TH staining in the SNpc and striatum. (M-N) Quantification of the number of TH-positive neurons and optical density of striatal TH expression. The number of TH-positive cells in SNpc and the optical density of TH-positive dopaminergic fibers in the striatum were significantly blocked by MCHR1 gene-silencing in the SNpc, but less by gene-silencing in the HPC (One-way ANOVA; for SNpc: $F_{5, 47} = 35.0$, $P < 0.001$; for striatum: $F_{5, 50} = 89.5$, $P < 0.001$; *post-hoc* Tukey's test: $^{***}P < 0.001$, $^{**}P < 0.01$). Data are shown as mean ± SEM.

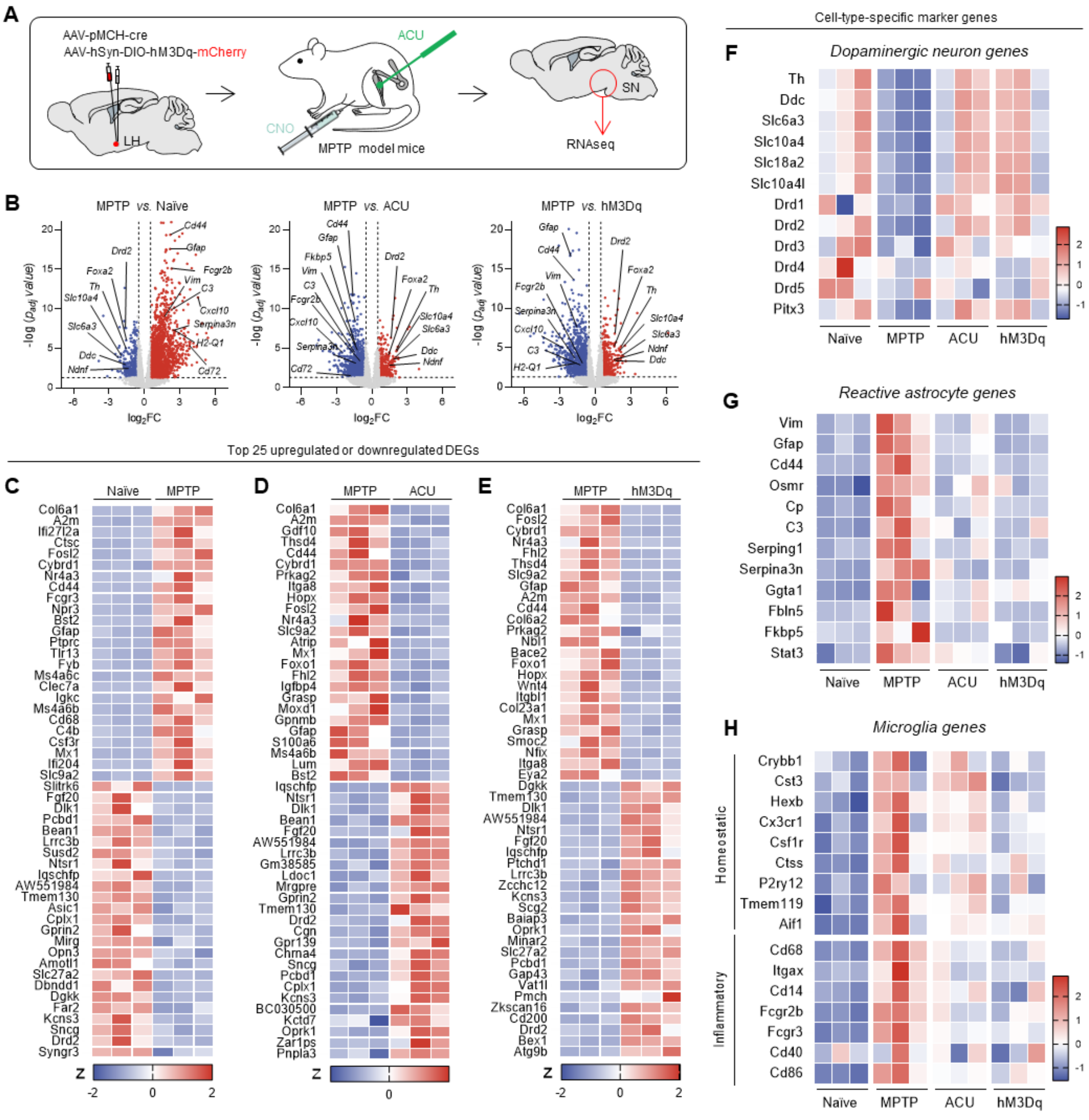


**Fig. S8. Transcriptomic change in the SNpc by acupuncture or chemogenetic activation of MCH neurons in MPTP model mice.** (A) Schematic diagram of SNpc tissue preparation for RNA-seq. (B) Volcano plot displaying the transcriptomic differences in the SNpc between MPTP and Naïve (left), between MPTP and ACU (middle), and MPTP and hM3Dq groups (right). Upregulated DEGs are marked in red, while downregulated DEGs are marked in blue. DEGs were defined with the criteria of $p_{adj} < 0.05$ and $|FC| > 1.5$. (C) Top 25 upregulated and downregulated SNpc DEGs by MPTP treatment with the highest statistical significance. DEGs were identified by three criteria: $p_{adj} < 0.05$, $|FC| \geq 1.5$, and FPKM (in any group) > 1.0. (D) Top 25 downregulated and upregulated SNpc DEGs by acupuncture treatment in MPTP model with the highest statistical significance. (E) Top 25 downregulated and upregulated SNpc DEGs by hM3Dq-mediated chemogenetic activation of MCH neurons in MPTP model with the highest statistical significance. (F-H) Heatmap showing enriched genes in specific cell types such as DA neurons (F), reactive astrocytes (G), and microglia (H) in the SNpc.

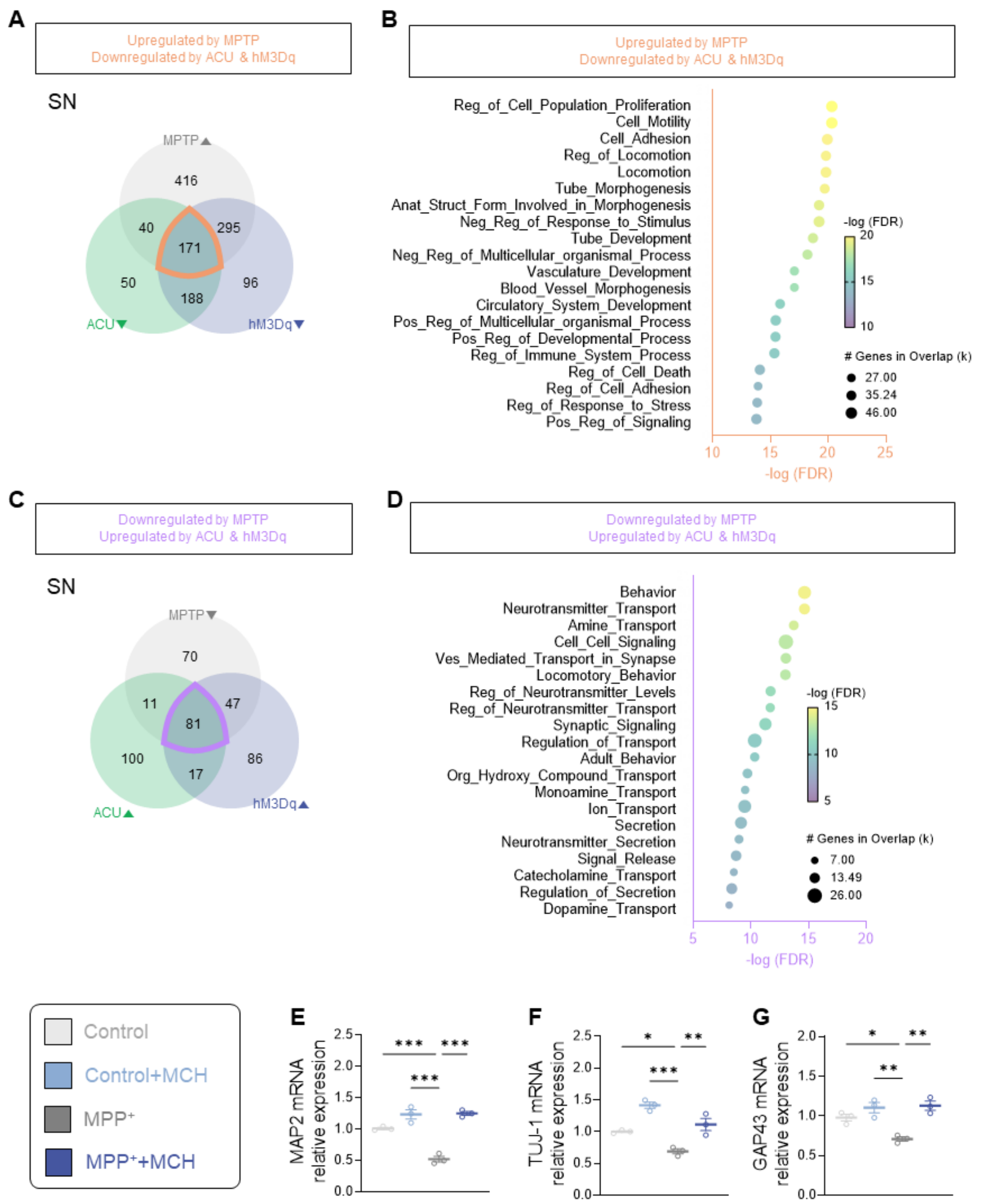


**Fig. S9. GO analysis of SNpc DEGs.** (A) Venn diagrams depicting the intersection of upregulated DEGs by MPTP (MPTP-up) and downregulated DEGs by either acupuncture or hM3Dq-mediated activation (ACU-down or hM3Dq-down). (B) Top 20 gene ontology (GO) terms analyzed from the intersection of MPTP-up, ACU-down, and hM3Dq-down (marked in orange in A) with the highest statistical significance. (C) Venn diagrams depicting the intersection of downregulated DEGs by MPTP (MPTP-down) and upregulated DEGs by either acupuncture or hM3Dq-mediated activation (ACU-up or hM3Dq-up). Note that many of the terms are related to the cellular reaction upon neuroinflammation. (D) Top 20 gene ontology (GO) terms analyzed from the intersection of MPTP-down, ACU-up, and hM3Dq-up (marked in violet in C) with the highest statistical significance. Note that many of the terms are related to the synthesis, transport, signaling, and secretion of amine neurotransmitter (especially, dopamine). (E-G) mRNA expressions of MAP2 (E), TUJ-1 (F), and GAP43 (G) assessed by RT-qPCR at 5 days after treatment of MCH along with $MPP^+$ (One-way ANOVA; for MAP2: $F_{3, 8} = 51.49$, $P <$

0.001; for TUJ-1: $F_{3, 8} = 26.63$, $P = 0.321$; for GAP43: $F_{3, 8} = 14.55$, $P = 0.8207$; *post-hoc* Tukey's test: $^{***}P < 0.001$, $^{**}P < 0.01$, $^{*}P < 0.05$). Data are shown as mean ± SEM.

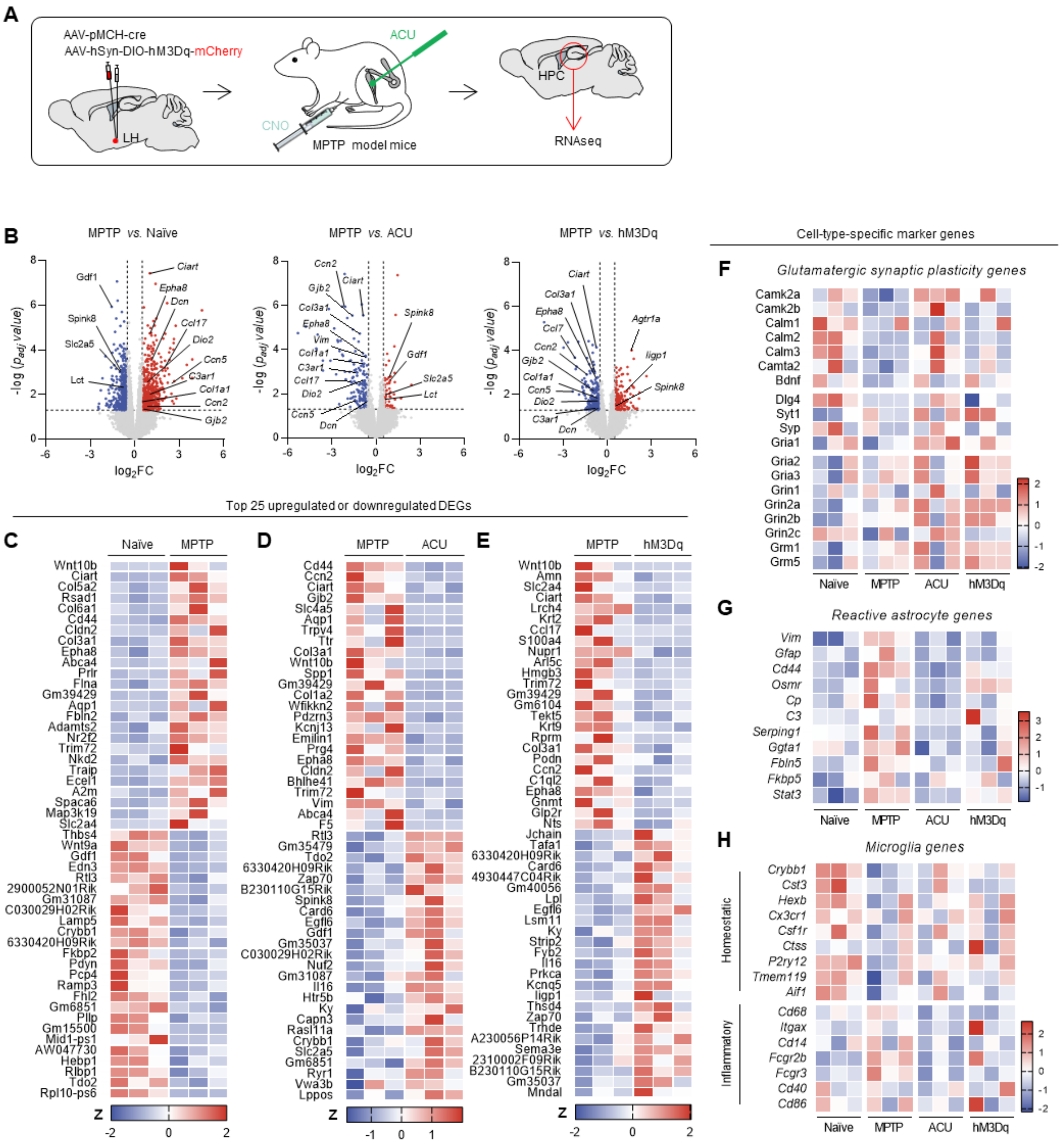


**Fig. S10. Transcriptomic change in the HPC by acupuncture or chemogenetic activation of MCH neurons in MPTP model mice.** (A) Schematic diagram of HPC tissue preparation for RNA-seq. (B) Volcano plot displaying the transcriptomic differences in the hippocampus between MPTP and Naïve (left), between MPTP 8 and ACU (middle), and MPTP and hM3Dq groups (right). DEGs were defined with the criteria of $p < 0.05$ and $|FC| > 1.5$. (C) Top 25 upregulated and downregulated hippocampal DEGs by MPTP treatment with the highest statistical significance. DEGs were identified by three criteria: $p < 0.05$, $|FC| \geq 1.5$, and FPKM (in any group) > 1.0. (D) Top 25 downregulated and upregulated hippocampal DEGs by acupuncture treatment in MPTP model with the highest statistical significance. (E) Top 25 downregulated and upregulated hippocampal DEGs by hM3Dq-mediated chemogenetic activation of MCH neurons in MPTP model with the highest statistical significance. (F-H) Heatmap showing enriched genes in specific cell types such as glutamatergic synapses (F), reactive astrocytes (G), and microglia (H).

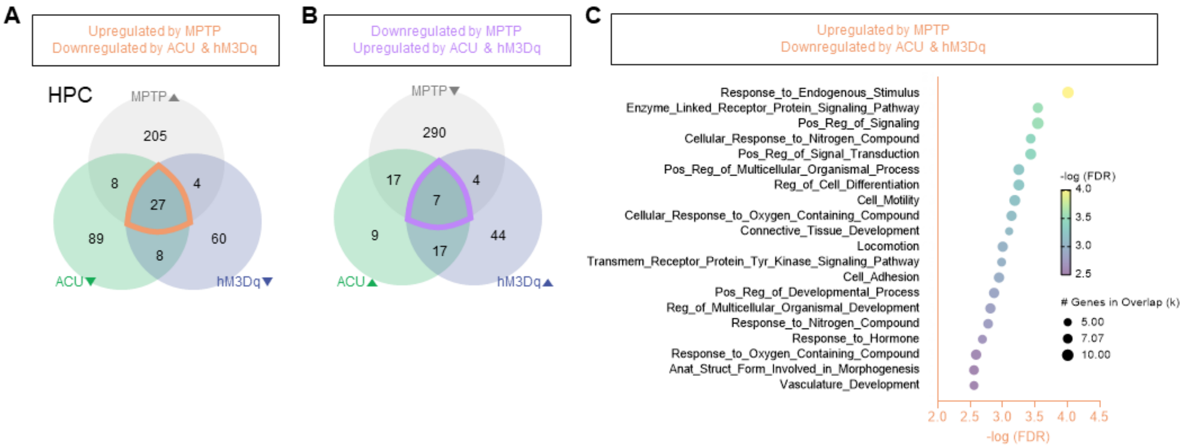


**Fig. S11. GO analysis of HPC DEGs.** (A) Venn diagrams depicting the intersection of upregulated DEGs by MPTP (MPTP-up) and downregulated DEGs by either acupuncture or hM3Dq-mediated activation (ACU-down or hM3Dq-down). (B) Venn diagrams depicting the intersection of downregulated DEGs by MPTP (MPTP-down) and upregulated DEGs by either acupuncture or hM3Dq-mediated activation (ACU-up or hM3Dq-up). (C) Top 20 gene ontology (GO) terms analyzed from the intersection of MPTP-up, ACU-down, and hM3Dq-down (marked in orange in H) with the highest statistical significance. Due to the small number of genes in the intersection of MPTP-down, ACU-up, and hM3Dq-up (marked in violet in I), GO analysis was not available.

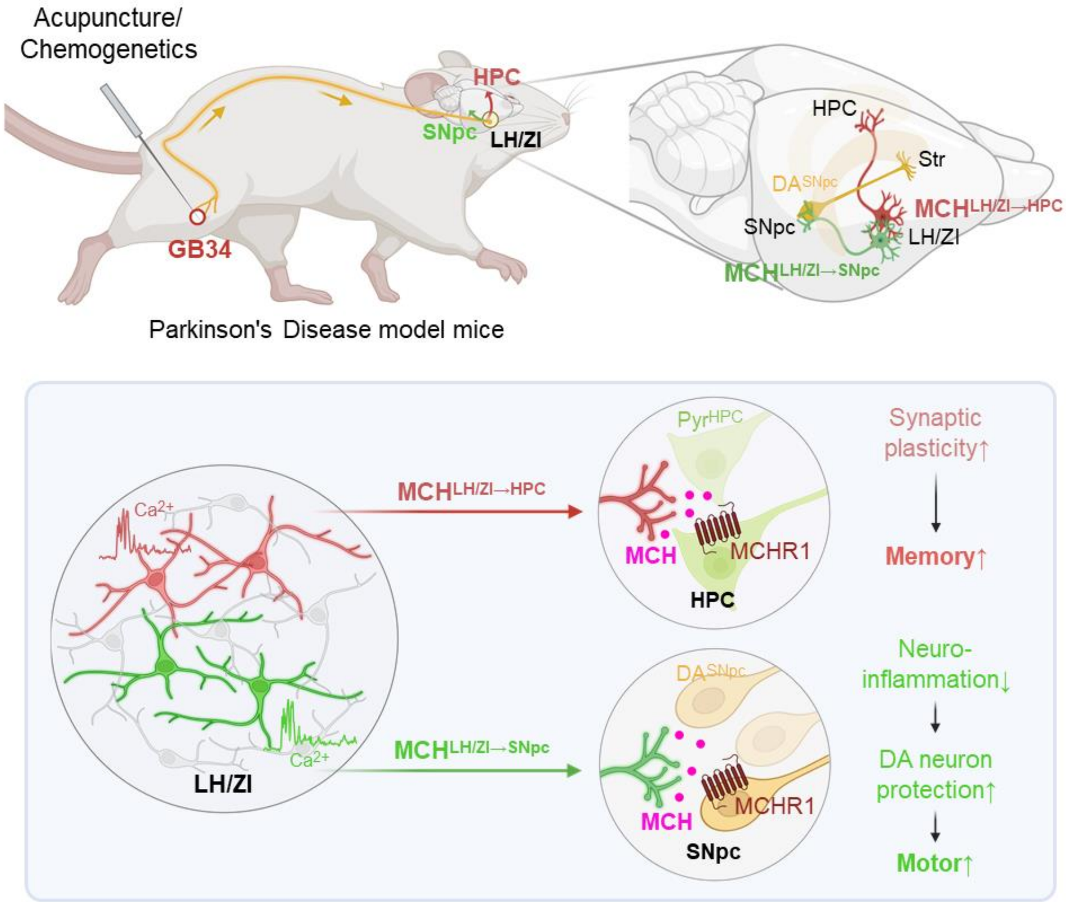


**Fig. S12. Schematic diagram of the mechanism underlying the acupuncture effect on motor and non-motor symptoms in the PD mouse model.** Acupuncture stimulation at a hindlimb acupoint GB34 activates the sensory afferents which are connected to the LH/ZI, leading to activation of MCH neurons in the LH and ZI ($MCH^{LH/ZI}$ neurons). Likewise, chemogenetic activation of the sensory afferents at GB34 activates the $MCH^{LH/ZI}$ neurons. $MCH^{LH/ZI}$ neurons project to HPC and SNpc which originate from distinct subpopulations. Activation of $MCH^{LH\rightarrow HPC}$ projections results in enhancement of hippocampal synaptic plasticity which causes memory improvement in the PD mouse model. On the other hand, activation of $MCH^{LH/ZI\rightarrow SNpc}$ projections exerts a strong anti-inflammatory effect through MCHR1-dependent manner, causing the protection of nigrostriatal DA neurons and improving the motor function in the PD mouse model. In summary, activations of $MCH^{LH\rightarrow HPC}$ and $MCH^{LH/ZI\rightarrow SNpc}$ projections are critical for alleviating the memory and motor dysfunction by acupuncture, respectively. Image was created with Biorender.com.

**Table S1. The detailed information about statistical analyses.**

| Figure No. | Result from statistical analysis |
|---|---|
| Figure 1F | Naïve (397 ± 8.3%, n = 16), MPTP (179.3 ± 17.0%, n = 16), MPTP+ACU (367.7 ± 13.1%, n = 16), MPTP+Lido+ACU (222.8 ± 17.2%, n = 12), SCNx+MPTP+ACU (39.2 ± 12.6%, n = 5), MPTP+nonACU (164.2 ± 10.8%, n = 10)<br>The data were not normally distributed.<br>Kruskal-Wallis ANOVA test with Dunn's multiple comparison test<br>$H (6) = 59.65$, $p < 0.001$ |
| Figure 1G | Naïve (24.5 ± 1.9%, n = 14), MPTP (6.7 ± 0.8%, n = 14), MPTP+ACU (21.7 ± 1.2%, n = 14), MPTP+Lido+ACU (8.3 ± 0.9%, n = 14), SCNx+MPTP+ACU (5.6 ± 1.0%, n = 8), MPTP+nonACU (5.9 ± 1.0%, n = 14)<br>One-way ANOVA with Tukey's multiple comparisons test<br>$F (5, 84) = 70.56$, $p < 0.001$ |
| Figure 1I | Naïve (9.5 ± 0.2%, n = 14), MPTP (5.4 ± 0.2%, n = 14), MPTP+ACU (8.8 ± 0.3%, n = 14), MPTP+Lido+ACU (5.7 ± 0.2%, n = 10), SCNx+MPTP+ACU (5.3 ± 0.2%, n = 10), MPTP+nonACU (5.5 ± 0.3%, n = 11)<br>One-way ANOVA with Tukey's multiple comparisons test<br>$F (5, 67) = 54.91$, $p < 0.001$ |
| Figure 1J | Naïve (39.0 ± 1.6%, n = 14), MPTP (19.5 ± 1.4%, n = 14), MPTP+ACU (33.2 ± 1.3%, n = 14), MPTP+Lido+ACU (22.6 ± 1.3%, n = 10), SCNx+MPTP+ACU (21.3 ± 1.3%, n = 10), MPTP+nonACU (19.1 ± 1.0%, n = 16)<br>One-way ANOVA with Tukey's multiple comparisons test<br>$F (5, 72) = 41.20$, $p < 0.001$ |
| Figure 1K | Naïve (72.5 ± 3.3%, n = 11), MPTP (49.3 ± 3.0%, n = 11), MPTP+ACU (69.5 ± 3.6%, n = 11), MPTP+Lido+ACU (41.3 ± 2.7%, n = 11), SCNx+MPTP+ACU (34.9 ± 3.7%, n = 11), MPTP+nonACU (44.9 ± 3.3%, n = 11)<br>One-way ANOVA with Tukey's multiple comparisons test<br>$F (5, 60) = 25.49$, $p < 0.001$ |
| Figure 1L | Naïve (73.0 ± 3.0%, n = 9), MPTP (47.7 ± 3.1%, n = 9), MPTP+ACU (71.8 ± 3.1%, n = 9), MPTP+Lido+ACU (45.9 ± 3.9%, n = 9), SCNx+MPTP+ACU (44.3 ± 2.7%, n = 9), MPTP+nonACU (43.8 ± 3.3%, n = 9)<br>One-way ANOVA with Tukey's multiple comparisons test<br>$F (5, 48) = 18.93$, $p < 0.001$ |
| Figure 1O | Naïve (414.2 ± 31.3%, n = 5), MPTP (106.1 ± 10.2%, n = 11), MPTP+CNO (302.2 ± 33.1%, n = 11)<br>One-way ANOVA with Tukey's multiple comparisons test<br>$F (2, 24) = 31.0$, $p < 0.001$ |
| Figure 1P | Naïve (23.4 ± 1.9%, n = 5), MPTP (5.0 ± 0.4%, n = 11), MPTP+CNO (14.2 ± 1.0%, n = 11)<br>One-way ANOVA with Tukey's multiple comparisons test<br>$F (2, 24) = 74.1$, $p < 0.001$ |
| Figure 1Q | Naïve (9.2 ± 0.3%, n = 5), MPTP (5.3 ± 0.4%, n = 11), MPTP+CNO (7.7 ± 0.3%, n = 11)<br>One-way ANOVA with Tukey's multiple comparisons test |

| | |
|---|---|
| | F (2, 24) = 26.9, p < 0.001 |
| Figure 1R | Naïve (33.0 ± 0.8%, n = 5), MPTP (15.7 ± 0.7%, n = 11), MPTP+CNO (23.1 ± 0.9%, n = 11)<br>One-way ANOVA with Tukey's multiple comparisons test<br>F (2, 24) = 84.1, p < 0.001 |
| Figure 1T | Naïve (75.6 ± 3.1%, n = 5), MPTP (45.4 ± 2.9%, n = 11), MPTP+CNO (68.0 ± 3.2%, n = 11)<br>One-way ANOVA with Tukey's multiple comparisons test<br>F (2, 24) = 22.8, p < 0.001 |
| Figure 1U | Naïve (74.7 ± 2.0%, n = 5), MPTP (40.6 ± 3.7%, n = 11), MPTP+CNO (56.5 ± 2.3%, n = 11)<br>One-way ANOVA with Tukey's multiple comparisons test<br>F (2, 24) = 22.6, p < 0.001 |
| Figure 2G | Rt. ACU (5.0 ± 0.5%, n = 11), Rt. nonACU (1.5 ± 0.4%, n = 11), Lido + Rt. AU (1.7 ± 0.4%, n = 11)<br>Repeated-Measure one-way ANOVA with Dunnett's multiple comparisons test (compared with Rt. ACU group)<br>F (2, 20) = 18.92, p < 0.001 |
| Figure 3B | Naïve (423.1 ± 13.0%, n = 19), MPTP (83.3 ± 13.6%, n = 15), MPTP+ACU (331.6 ± 19.0%, n = 9), MPTP+hM4Di+ACU (61.2 ± 7.6%, n = 13), MPTP+hM3Dq (355.2 ± 18.4%, n = 21)<br>One-way ANOVA with Tukey's multiple comparisons test<br>F (4, 72) = 112.8, p < 0.001 |
| Figure 3C | Naïve (19.8 ± 1.2%, n = 19), MPTP (3.8 ± 0.8%, n = 15), MPTP+ACU (21.1 ± 1.4%, n = 9), MPTP+hM4Di+ACU (3.9 ± 0.4%, n = 13), MPTP+hM3Dq (22.7 ± 1.8%, n = 21)<br>One-way ANOVA with Tukey's multiple comparisons test<br>F (4, 72) = 47.05, p < 0.001 |
| Figure 3e | Naïve (8.5 ± 0.3%, n = 11), MPTP (5.2 ± 0.3%, n = 11), MPTP+ACU (8.6 ± 0.2%, n = 11), MPTP+hM4Di+ACU (5.0 ± 0.2%, n = 11), MPTP+hM3Dq (8.4 ± 0.5%, n = 11)<br>The data were not normally distributed.<br>Kruskal-Wallis ANOVA test with a Dunn's multiple comparison test, p < 0.001<br>H (5) = 37.32, p < 0.001 |
| Figure 3F | Naïve (32.7 ± 0.5%, n = 10), MPTP (13.4 ± 0.5%, n = 15), MPTP+ACU (26.9 ± 0.8%, n = 12), MPTP+hM4Di+ACU (13.3 ± 0.7%, n = 12), MPTP+hM3Dq (28.7 ± 0.9%, n = 10)<br>One-way ANOVA with Tukey's multiple comparisons test<br>F (4, 54) = 177, p < 0.001 |
| Figure 3G | Naïve (75.1 ± 2.1%, n = 19), MPTP (42.7 ± 3.0%, n = 15), MPTP+ACU (75.6 ± 3.4%, n = 9), MPTP+hM4Di+ACU (34.3 ± 2.1%, n = 13), MPTP+hM3Dq (74.0 ± 1.8%, n = 21)<br>One-way ANOVA with Tukey's multiple comparisons test<br>F (4, 72) = 73.2, p < 0.001 |

| | |
|---|---|
| Figure 3H | Naïve (74.7 ± 1.3%, n = 19), MPTP (35.3 ± 3.3%, n = 17), MPTP+ACU (72.3 ± 1.7%, n = 9), MPTP+hM4Di+ACU (33.9 ± 3.3%, n = 15), MPTP+hM3Dq (69.9 ± 1.7%, n = 23)<br>One-way ANOVA with Tukey's multiple comparisons test<br>$F (4, 78) = 72.45$, $p < 0.001$ |
| Figure 3J | Naïve (168.7 ± 12.9%, n = 10), MPTP (101.8 ± 9.2%, n = 10), MPTP+ACU (183.7 ± 22.4%, n = 10), MPTP+hM4Di+ACU (99.2 ± 16.2%, n = 10), MPTP+hM3Dq (175.9 ± 25.6%, n = 10)<br>One-way ANOVA with Tukey's multiple comparisons test<br>$F (4, 45) = 5.213$, $p = 0.0015$ |
| Figure 4h | $mCherry^+$ (-33.4 ± 1.6%, n = 10), $GFP^+$ (-28.9 ± 0.8%, n = 10)<br>Unpaired two-tailed t-test, p = 0.023 |
| Figure 4I | Multiple unpaired two-tailed t-test<br>-120 pA: $mCherry^+$ (0.0 ± 0.0%, n = 10), $GFP^+$ (0.0 ± 0.0%, n = 10), p = 0.135<br>-100 pA: $mCherry^+$ (0.0 ± 0.0%, n = 10), $GFP^+$ (0.0 ± 0.0%, n = 10), p = 0.135<br>-80 pA: $mCherry^+$ (0.0 ± 0.0%, n = 10), $GFP^+$ (0.0 ± 0.0%, n = 10), p = 0.135<br>-60 pA: $mCherry^+$ (0.0 ± 0.0%, n = 10), $GFP^+$ (0.0 ± 0.0%, n = 10), p = 0.135<br>-40 pA: $mCherry^+$ (0.0 ± 0.0%, n = 10), $GFP^+$ (0.0 ± 0.0%, n = 10), p = 0.135<br>-20 pA: $mCherry^+$ (0.0 ± 0.0%, n = 10), $GFP^+$ (0.0 ± 0.0%, n = 10), p = 0.135<br>0 pA: $mCherry^+$ (0.6 ± 0.4%, n = 10), $GFP^+$ (0.1 ± 0.1%, n = 10), p = 0.269<br>20 pA: $mCherry^+$ (6.2 ± 2.3%, n = 10), $GFP^+$ (2.3 ± 1.1%, n = 10), p = 0.141<br>40 pA: $mCherry^+$ (10.9 ± 3.9%, n = 10), $GFP^+$ (5.6 ± 1.8%, n = 10), p = 0.233<br>60 pA: $mCherry^+$ (15.2 ± 5.2%, n = 10), $GFP^+$ (8.7 ± 2.9%, n = 10), p = 0.294<br>80 pA: $mCherry^+$ (18.4 ± 5.6%, n = 10), $GFP^+$ (10.7 ± 3.8%, n = 10), p = 0.271<br>100 pA: $mCherry^+$ (21.7 ± 5.8%, n = 10), $GFP^+$ (12.6 ± 4.5%, n = 10), p = 0.230<br>120 pA: $mCherry^+$ (21.9 ± 5.4%, n = 10), $GFP^+$ (13.8 ± 5.0%, n = 10), p = 0.286<br>140 pA: $mCherry^+$ (24.5 ± 5.6%, n = 10), $GFP^+$ (14.3 ± 5.0%, n = 10), p = 0.190<br>160 pA: $mCherry^+$ (26.4 ± 5.8%, n = 10), $GFP^+$ (11.7 ± 4.1%, n = 10), p = 0.054<br>180 pA: $mCherry^+$ (27.3 ± 6.0%, n = 10), $GFP^+$ (8.4 ± 2.9%, n = 10), p = 0.010<br>200 pA: $mCherry^+$ (29.3 ± 6.5%, n = 10), $GFP^+$ (6.7 ± 2.3%, n = 10), p = 0.004<br>220 pA: $mCherry^+$ (30.4 ± 7.0%, n = 10), $GFP^+$ (5.6 ± 1.7%, n = 10), p = 0.003 |
| Figure 4J | $mCherry^+$ (-120, 3.1 ± 0.8%, n = 10; -100, 3.0 ± 1.0%, n = 10; -80, 2.9 ± 1.0%, n = 10; -60, 2.7 ± 1.0%, n = 10; -40, 2.0 ± 1.1%, n = 10; -20, 1.4 ± 1.0%, n = 10; 0, 0.1 ± 0.1%, n = 10; 20, 0.0 ± 0.0%, n = 10; 40, 0.0 ± 0.0%, n = 10; 60, 0.0 ± 0.0%, n = 10; 80, 0.0 ± 0.0%, n = 10; 100, 0.0 ± 0.0%, n = 10; 120, 0.0 ± 0.0%, n = 10; 140, 0.0 ± 0.0%, n = 10; 160, 0.0 ± 0.0%, n = 10; 180, 0.0 ± 0.0%, n = 10; 200, 0.0 ± 0.0%, n = 10; 220, 0.0 ± 0.0%, n = 10), $GFP^+$ (-120, 1.1 ± 0.4%, n = 10; -100, 0.9 ± 0.3%, n = 10; -80, 0.8 ± 0.3%, n = 10; -60, 0.5 ± 0.2%, n = 10; -40, 0.1 ± 0.1%, n = 10; -20, 0.1 ± 0.1%, n = 10; 0, 0.0 ± 0.0%, n = 10; 20, 0.0 ± 0.0%, n = 10; 40, 0.0 ± 0.0%, n = 10; 60, 0.0 ± 0.0%, n = 10; 80, 0.0 ± 0.0%, n = 10; 100, 0.0 ± 0.0%, n = 10; 120, 0.0 ± 0.0%, n = 10; 140, 0.0 ± 0.0%, n = 10; 160, 0.0 ± 0.0%, n = 10; 180, 0.0 ± 0.0%, n = 10; 200, 0.0 ± 0.0%, n = 10; 220, 0.0 ± 0.0%, n = 10)<br>Two-way ANOVA with Sidak's multiple comparisons test<br>Interaction, $F (17, 306) = 3.604$, $p < 0.0001$; Injected currents, $F (1.235, 22.22) = 10.33$, $p = 0.0025$; Neuron subtype, $F (1, 18) = 3.997$, $p = 0.0609$ |

| | |
|---|---|
| Figure 4L | Naïve (398.7 ± 19.0%, n = 13), MPTP (25.2 ± 3.9%, n = 14), MPTP+ACU (370.2 ± 18.7%, n = 11), MPTP+hM4Di+ACU (48.4 ± 7.5%, n = 9), MPTP+hM3Dq (215.9 ± 16.8%, n = 8)<br>One-way ANOVA with Tukey's multiple comparisons test<br>$F (4, 50) = 156, p < 0.001$ |
| Figure 4M | Naïve (17.9 ± 1.5%, n = 16), MPTP (4.8 ± 0.3%, n = 14), MPTP+ACU (17.2 ± 0.5%, n = 11), MPTP+hM4Di+ACU (6.6 ± 0.6%, n = 9), MPTP+hM3Dq (18.8 ± 0.9%, n = 8)<br>One-way ANOVA with Tukey's multiple comparisons test<br>$F (4, 53) = 46.08, p < 0.001$ |
| Figure 4N | Naïve (79.0 ± 1.9%, n = 16), MPTP (35.1 ± 3.0%, n = 13), MPTP+ACU (80.6 ± 2.1%, n = 11), MPTP+hM4Di+ACU (66.8 ± 2.9%, n = 9), MPTP+hM3Dq (34.3 ± 3.7%, n = 8)<br>One-way ANOVA with Tukey's multiple comparisons test<br>$F (4, 52) = 72.8, p < 0.001$ |
| Figure 4O | Naïve (70.2 ± 2.4%, n = 16), MPTP (35.5 ± 3.1%, n = 19), MPTP+ACU (69.5 ± 2.4%, n = 11), MPTP+hM4Di+ACU (57.5 ± 1.2%, n = 9), MPTP+hM3Dq (38.1 ± 2.5%, n = 8)<br>One-way ANOVA with Tukey's multiple comparisons test<br>$F (4, 58) = 39.11, p < 0.001$ |
| Figure 4Q | Naïve (384.3 ± 19.0%, n = 16), MPTP (61.6 ± 12.5%, n = 14), MPTP+ACU (336.0 ± 24.9%, n = 11), MPTP+hM4Di+ACU (199.4 ± 17.9%, n = 10), MPTP+hM3Dq (45.4 ± 8.0%, n = 10)<br>One-way ANOVA with Tukey's multiple comparisons test<br>$F (4, 56) = 79.68, p < 0.001$ |
| Figure 4R | Naïve (20.3 ± 0.8%, n = 16), MPTP (4.6 ± 0.5%, n = 14), MPTP+ACU (17.5 ± 0.7%, n = 11), MPTP+hM4Di+ACU (14.2 ± 0.6%, n = 10), MPTP+hM3Dq (8.5 ± 1.2%, n = 10)<br>One-way ANOVA with Tukey's multiple comparisons test<br>$F (4, 56) = 82.8, p < 0.001$ |
| Figure 4S | Naïve (76.3 ± 2.6%, n = 16), MPTP (34.1 ± 1.8%, n = 14), MPTP+ACU (77.6 ± 2.0%, n = 11), MPTP+hM4Di+ACU (44.8 ± 2.4%, n = 10), MPTP+hM3Dq (67.4 ± 4.1%, n = 10)<br>One-way ANOVA with Tukey's multiple comparisons test<br>$F (4, 56) = 60.4, p < 0.001$ |
| Figure 4T | Naïve (73.3 ± 1.3%, n = 16), MPTP (38.1 ± 3.1%, n = 15), MPTP+ACU (74.2 ± 1.5%, n = 11), MPTP+hM4Di+ACU (39.8 ± 2.1%, n = 10), MPTP+hM3Dq (70.0 ± 2.4%, n = 10)<br>One-way ANOVA with Tukey's multiple comparisons test<br>$F (4, 57) = 67.9, p < 0.001$ |
| Figure 5B | Naïve (430.8 ± 18.2%, n = 6), MPTP (142.7 ± 15.8%, n = 6), MPTP+ACU (355.7 ± 25.2%, n = 6), MPTP+MCH (274.7 ± 52.2%, n = 6), MPTP+7c+ACU (182.5 ± 26.2%, n = 6), MPTP+7c (166.8 ± 29.5%, n = 6)<br>One-way ANOVA with Tukey's multiple comparisons test<br>$F (5, 30) = 14.6, p < 0.001$ |

| | |
|---|---|
| Figure 5C | Naïve (14.7 ± 1.2%, n = 6), MPTP (3.0 ± 1.0%, n = 6), MPTP+ACU (12.3 ± 1.0%, n = 6), MPTP+MCH (9.0 ± 1.4%, n = 6), MPTP+7c+ACU (6.2 ± 1.0%, n = 6), MPTP+7c (3.0 ± 0.7%, n = 6)<br>One-way ANOVA with Tukey's multiple comparisons test<br>$F(5, 30) = 19.8$, $p < 0.001$ |
| Figure 5D | Naïve (75.4 ± 3.8%, n = 6), MPTP (50.5 ± 2.1%, n = 6), MPTP+ACU (70.8 ± 1.7%, n = 6), MPTP+MCH (62.2 ± 2.5%, n = 6), MPTP+7c+ACU (52.5 ± 3.3%, n = 6), MPTP+7c (46.8 ± 6.9%, n = 6)<br>The data were not normally distributed.<br>Kruskal-Wallis ANOVA test with a Dunn's multiple comparison test, $p < 0.001$<br>$H(6) = 25.46$, $p < 0.001$ |
| Figure 5E | Naïve (75.7 ± 3.4%, n = 6), MPTP (42.3 ± 5.3%, n = 6), MPTP+ACU (70.0 ± 2.5%, n = 6), MPTP+MCH (65.6 ± 3.0%, n = 6), MPTP+7c+ACU (51.6 ± 2.0%, n = 6), MPTP+7c (45.9 ± 4.2%, n = 6)<br>The data were not normally distributed.<br>Kruskal-Wallis ANOVA test with a Dunn's multiple comparison test, $p < 0.001$<br>$H(6) = 28.31$, $p < 0.001$ |
| Figure 5G | Naïve (361.9 ± 11.5%, n = 13), MPTP (96.3 ± 7.6%, n = 16), MPTP+ACU+shScr (SNpc) (321.6 ± 18.9%, n = 8), MPTP+ACU+shMCHR1 (SNpc) (189.4 ± 21.9%, n = 8), MPTP+ACU+shScr (HPC) (350.4 ± 14.6%, n = 8), MPTP+ACU+shMCHR1 (HPC) (339.0 ± 13.8%, n = 8)<br>One-way ANOVA with Tukey's multiple comparisons test<br>$F(5, 55) = 79.2$, $p < 0.001$ |
| Figure 5H | Naïve (25.2 ± 1.1%, n = 13), MPTP (11.4 ± 0.8%, n = 15), MPTP+ACU+shScr (SNpc) (22.8 ± 1.0%, n = 9), MPTP+ACU+shMCHR1 (SNpc) (16.4 ± 1.1%, n = 9), MPTP+ACU+shScr (HPC) (26.1 ± 0.9%, n = 8), MPTP+ACU+shMCHR1 (HPC) (24.4 ± 1.4%, n = 8)<br>One-way ANOVA with Tukey's multiple comparisons test<br>$F(5, 56) = 38.2$, $p < 0.001$ |
| Figure 5I | Naïve (78.5 ± 1.5%, n = 13), MPTP (57.3 ± 1.8%, n = 16), MPTP+ACU+shScr (SNpc) (71.5 ± 2.7%, n = 8), MPTP+ACU+shMCHR1 (SNpc) (62.4 ± 4.5%, n = 9), MPTP+ACU+shScr (HPC) (75.2 ± 1.9%, n = 8), MPTP+ACU+shMCHR1 (HPC) (59.1 ± 3.8%, n = 8)<br>The data were not normally distributed.<br>Kruskal-Wallis ANOVA test with a Dunn's multiple comparison test, $p < 0.001$<br>$H(6) = 37.75$, $p < 0.001$ |
| Figure 5J | Naïve (79.6 ± 3.1%, n = 13), MPTP (49.5 ± 3.1%, n = 16), MPTP+ACU+shScr (SNpc) (73.4 ± 5.8%, n = 8), MPTP+ACU+shMCHR1 (SNpc) (60.7 ± 4.3%, n = 9), MPTP+ACU+shScr (HPC) (77.3 ± 2.8%, n = 8), MPTP+ACU+shMCHR1 (HPC) (56.5 ± 2.4%, n = 8)<br>One-way ANOVA with Tukey's multiple comparisons test<br>$F(5, 56) = 14.2$, $p < 0.001$ |
| Figure 6E | Control (90.3 ± 4.4%, n = 6), $MPP^+$ (23.8 ± 3.9%, n = 6), Control+MCH (91.8 ± 5.4%, n = 6), $MPP^+$+MCH (54.8 ± 4.2%, n = 6)<br>One-way ANOVA with Tukey's multiple comparisons test<br>$F(3, 20) = 51.42$, $p < 0.001$ |

| | |
|---|---|
| Figure 6F | Control (98.0 ± 3.8%, n = 6), $MPP^+$ (23.0 ± 3.4%, n = 6), Control+MCH (95.3 ± 6.2%, n = 6), $MPP^+$+MCH (55.3 ± 3.8%, n = 6)<br>One-way ANOVA with Tukey's multiple comparisons test<br>F (3, 20) = 65.05, p < 0.001 |
| Figure 6G | Control (0.9 ± 0.0%, n = 3), $MPP^+$ (0.7 ± 0.0%, n = 3), Control+MCH (1.1 ± 0.0%, n = 3), $MPP^+$+MCH (1.0 ± 0.0%, n = 3)<br>One-way ANOVA with Tukey's multiple comparisons test<br>F (3, 8) = 39.27, p < 0.001 |
| Figure 6I | Naïve (760.5 ± 43.3%, n = 85), MPTP (1457.6 ± 209.4%, n = 23), MPTP+ACU (467.2 ± 23.4%, n = 89), MPTP+hM3Dq (582.7 ± 41.8%, n = 44)<br>One-way ANOVA with Tukey's multiple comparisons test<br>F (3, 237) = 34.6, p < 0.001 |
| Figure 6J | Naïve (125.0 ± 12.3%, n = 79), MPTP (550.1 ± 91.7%, n = 23), MPTP+ACU (121.9 ± 10.6%, n = 89), MPTP+hM3Dq (223.6 ± 25.3%, n = 44)<br>One-way ANOVA with Tukey's multiple comparisons test<br>F (3, 231) = 39.82, p < 0.001 |
| Figure 6K | Naïve (474.7 ± 30.9%, n = 42), MPTP (885.4 ± 54.2%, n = 39), MPTP+ACU (631.5 ± 35.7%, n = 46), MPTP+hM3Dq (622.9 ± 34.1%, n = 42)<br>The data were not normally distributed.<br>Kruskal-Wallis ANOVA test with Dunn's multiple comparison test<br>H (4) = 40.60, p < 0.001 |
| Figure 6L | Naïve (82.8 ± 8.1%, n = 42), MPTP (320.2 ± 18.8%, n = 39), MPTP+ACU (125.2 ± 9.7%, n = 100), MPTP+hM3Dq (206.6 ± 17.9%, n = 42)<br>The data were not normally distributed.<br>Kruskal-Wallis ANOVA test with Dunn's multiple comparison test<br>H (4) = 83.21, p < 0.001 |
| Figure 6N | Two-way ANOVA with Tukey's multiple comparisons test<br>Interaction, F (42,1935) = 7.716, p < 0.001; Radius, F (14, 1935) = 328.4, p < 0.001; Group, F (3, 1935) = 110.6, p < 0.001 |
| Figure 6O | Naïve (19.7 ± 1.3%, n = 26), MPTP (40.3 ± 2.8%, n = 34), MPTP+ACU (23.0 ± 1.3%, n = 36), MPTP+hM3Dq (20.1 ± 1.1%, n = 40)<br>The data were not normally distributed.<br>Kruskal-Wallis ANOVA test with Dunn's multiple comparison test<br>H (4) = 52.46, p < 0.001 |
| Figure 6P | Naïve (2.4 ± 0.8%, n = 28), MPTP (5.8 ± 2.9%, n = 36), MPTP+ACU (2.6 ± 0.9%, n = 38), MPTP+hM3Dq (2.9 ± 1.2%, n = 42)<br>The data were not normally distributed.<br>Kruskal-Wallis ANOVA test with Dunn's multiple comparison test<br>H (4) = 20.82, p < 0.001 |
| Figure 6T | Naïve (138.8 ± 12.2%, n = 5), $MPP^+$ (95.1 ± 7.4%, n = 5), MPTP+MCH100 (127.2 ± 19.7%, n = 5), MPTP+MCH200 (158.1 ± 18.3%, n = 5)<br>One-way ANOVA with Tukey's multiple comparisons test<br>F (3, 16) = 3.016, p = 0.0607 |

**Table S2. The detailed information about virus titers, volumes, and total amount for each experiment.**

| Figure | Virus | Titer (GC/ml) | Volume (μL) | Total amount (GC) |
|---|---|---|---|---|
| 1M-U, 2A-G, 4A-J, 6A-C, 6H-Q, S3A-B, S3L-N, S4A-B, S8A-H, S9A-D, S10A-H, S11A-C | $AAV_{DJ}$-pMCH-cre | 5.7x10^13 | 1.5 | 8.55x10^10 |
| 2A-C, S2E-H | PRV-CAG-EGFP | 3.0x10^9 | 1.5 | 4.5x10^6 |
| S2F-H | PRV-CAG-RFP | 3.0x10^9 | 1.5 | 4.5x10^6 |
| 4A-B, S4A-B | $AAV_{DJ}$-hSyn-EGFP | 9.17x10^12 | 1.5 | 1.38x10^10 |
| 4C-J | $AAV_{retro}$-hSyn-DIO-mCherry | 2.23x10^13 | 1.5 | 3.35x10^10 |
| 4C-J | $AAV_{retro}$-hSyn-DIO-EGFP | 1.8x10^13 | 1.5 | 2.7x10^10 |
| S2A, S3C-L, S4C, S5A-H | AAV-pMCH-EGFP-cre | 5.96x10^13 | 1.5 | 8.94x10^10 |
| 3A-J, 4K-T, 6A-C, 6H-Q, S3B-N, S4D-J, S5A-H, S6A-H, S8A-H, S9A-D, S10A-H, S11A-C | $AAV_{DJ}$-hSyn-DIO-hM3Dq-mCherry | 8.77x10^12 | 1.5 | 1.32x10^10 |
| S5A-D | AAV-DDC-cre | 8.9x10^13 | 1.5 | 1.34x10^11 |
| 3A-J, 4K-T, S3A, S3L-N, S6A-H | $AAV_{DJ}$-hSyn-DIO-hM4Di-mCherry | 1.66x10^13 | 1.5 | 2.49x10^10 |
| 4K-T, S4D-I, S6A-H | $AAV_{retro}$-pMCH-EGFP-cre | 8.71x10^13 | 1.5 | 1.31x10^11 |
| 5F-J, S7J-N | $AAV_{DJ}$-pSicoR-MCHR1sh-mCherry | 2.31x10^13 | 1.5 | 3.47x10^10 |
| 5F-J, S7J-N | $AAV_{DJ}$-pSicoR-scr-shRNA-mCherry | 4.8x10^13 | 1.5 | 7.2x10^10 |
| 1M-U, S2B-D | $AAV_{retro}$-hSyn-hM3Dq-mCherry | 3.53x10^13 | 1.5 | 5.3x10^10 |
| S3C-I | $AAV_{DJ}$-CMV-A53T-SNCA | 2.42x10^13 | 1.5 | 3.63x10^10 |

**Table S3. Sequences of oligonucleotides**

| | |
|---|---|
| shRNA targeting sequence: Mchr1 #1 | 5'-GCA CAA GGA GTG TCT CCT ACA-3' |
| shRNA targeting sequence: Mchr1 #2 | 5'-GCA ACG TCC CTG ACA TCT TCA-3' |
| shRNA targeting sequence: Mchr1 #3 | 5'-GCC TCA ATC CCT TTG TGT ACA-3' |
| qPCR primer for TH Forward | AGG TCT ACA CCA CGC TGA AG |
| qPCR primer for TH Reverse | TAC TGG GTG CAC TGG AAC AC |
| qPCR primer for MAP2 Forward | CTG GCA CCC CAC CAA GTT AT |
| qPCR primer for MAP2 Reverse | CTT CAG GTC TGG CAG TGG TT |
| qPCR primer for GAP43 Forward | CCG ATG GGG TGG AGA AGA AG |
| qPCR primer for GAP43 Reverse | GGA GGA CGG CGA GTT ATC AG |
| qPCR primer for TUJ1 Forward | CAA CGA GGC CTC TTC TCA CA |
| qPCR primer for TUJ1 Reverse | CAG GCA GTC GCA GTT TTC AC |
